# Single Nuclei Sequencing of Human Putamen Oligodendrocytes Reveals Altered Heterogeneity and Disease-Associated Changes in Parkinson’s Disease and Multiple System Atrophy

**DOI:** 10.1101/2021.05.06.442967

**Authors:** Erin Teeple, Pooja Joshi, Rahul Pande, Yinyin Huang, Akshat Karambe, Martine Latta-Mahieu, S. Pablo Sardi, Angel Cedazo-Minguez, Katherine W. Klinger, Amilcar Flores-Morales, Stephen L. Madden, Deepak Rajpal, Dinesh Kumar

## Abstract

The role of oligodendrocytes in neurodegenerative diseases remains incompletely understood and largely unexplored at the single cell level. We profiled 87,086 single nuclei from human brain putamen region for healthy control, Parkinson’s Disease (PD), and Multiple System Atrophy (MSA). Oligodendrocyte lineage cells were the dominant cell-type in the putamen with oligodendrocyte subpopulations clustered by transcriptomic variation found to exhibit diverse functional enrichment patterns, and this oligodendrocyte heterogeneity was altered in a disease-specific way. Among profiled oligodendrocyte subpopulations, differences in expression of SNCA, HAPLN2, MAPT, APP, and OPALIN were observed for PD and MSA compared with healthy controls. Intriguingly, greater activation of unfolded protein response pathway gene expression was observed in PD nuclei versus MSA. Using network analysis, we then identified specific PD- and MSA-correlated gene co-expression modules enriched with disease relevant pathways; the PD-correlated module was significantly enriched for Parkinson’s Disease GWAS loci (p = 0.01046). Our analysis provides a broader understanding of oligodendrocyte heterogeneity and reveals distinctive oligodendrocyte pathological alterations associated with PD and MSA which may suggest potential novel therapeutic targets and new strategies for disease modification.

## Introduction

Parkinson’s Disease (PD) and Multiple System Atrophy (MSA) are each synucleinopathies, progressive neurodegenerative disorders characterized by nervous system aggregates of α-synuclein protein [1, 2]. PD is the most frequently occurring synucleinopathy and second most common chronic neurodegenerative disorder after Alzheimer’s Disease, with a median age-standardized incidence rate in high-income countries of 160 per 100,000 individuals over age 65 [1]. In contrast, MSA occurs at a much lower frequency than PD, with an estimated mean incidence rate of 0.6-3 per 100,000 people [2, 3]. In both PD and MSA, intracytoplasmic inclusions of α-synuclein are observed on post-mortem histologic examination, although the predominant cellular localization of α-synuclein differs between the two conditions: α-synuclein aggregates are observed mainly as neuronal intracellular aggregates (Lewy bodies) in PD [4, 5] while in MSA, these aggregates occur most frequently as oligodendroglial cytoplasmic inclusions [5, 6].

Recent genome-wide association studies (GWAS) including large PD patient cohorts and new results from single-nucleus transcriptomic profiling in midbrain samples have highlighted potential roles for glial cells in the synucleinopathies, with particularly strong evidence implicating oligodendrocytes in PD [3–7]. Oligodendrocyte lineage cells have been proposed to participate in synucleinopathy development and progression via multiple mechanisms including insufficient metabolic support, initiation and propagation of exaggerated stress and inflammatory response signaling, and altered autophagy functions [3–6]. In a recently published single-nucleus RNA-seq analysis of substantia nigra tissue from healthy donors, oligodendrocyte-specific differentially expressed genes were found to be associated with genes linked with variants significantly associated with PD by GWAS [3]. Consistent with these findings, Bryois et al. also reported significant associations between PD risk variants and oligodendrocyte cells when integrating GWAS data with mouse nervous system single-cell data [6]. Alpha-synuclein is a 140-amino acid protein which participates in diverse functions including vesicle exocytosis, endocytosis, and neurotransmitter vesicle cycling [8–10], and it has also been found to localize to the nucleus, where direct interaction with DNA and histone proteins [11, 12] and modulation of DNA damage responses [13] have been described. Yet which SNCA-linked functions are dysregulated in nervous system cell subpopulations remains incompletely understood among different synucleinopathies.

In this analysis, we explore how midbrain oligodendrocyte lineage cell populations may be involved in PD and MSA using single-nucleus RNA sequencing technologies. We found oligodendrocytes to be the dominant cell type in all putamen samples, comprising on average 70%, 66%, 68% of sample nuclei for Control, PD, and MSA samples. Among oligodendrocyte nuclei clustering by variable gene expression, functions including myelination, focal adhesion, and vesicle transport were found to be differentially enriched in these identified clusters. Within clustered subpopulations, differential gene expression and WGCNA analysis revealed that in comparison to Control, both PD and MSA oligodendrocyte nuclei exhibited distinct and differentiated transcriptional changes related to proteotoxic stress response, myelination, membrane process development, vesicular transport, and cell adhesion. Interestingly, SNCA expression was observed to increase most dramatically among oligodendrocyte clusters expressing greater levels of HAPLN2 specifically in PD. Integration of putamen data with data from healthy human substantia nigra samples then revealed that the oligodendrocyte subpopulation exhibiting the highest expression of HAPLN2 were predicted to comprise a significantly greater proportion of oligodendrocyte nuclei in substantia nigra compared with putamen (X-squared p < 2.2e-16). These results suggest that oligodendrocyte functional and population heterogeneity may be contributing factors in selective regional involvement in PD and possibly MSA. Our observation of distinctive patterns of proteotoxic stress response activation and differences in oligodendrocyte transcriptional activity for PD versus MSA suggest different pathological mechanisms for these SNCA-linked disorders with respect to oligodendrocytes.

## Results

### Oligodendrocytes Are the Major Cell Type Found in the Putamen

Single nuclei sequencing technologies were used to analyze human putamen tissue samples obtained postmortem from Control (n = 3 donors; 21,418 nuclei), PD (n=3 donors; 32,301 nuclei), and MSA (n = 3 donors; 32,488 nuclei) patients. Figure 1A presents an overall workflow schematic for the analysis. In order to first assess overall broad cell types proportions and to identify oligodendrocyte lineage nuclei, sample nuclei-gene count matrices were integrated and then clustered using the Seurat package v3.1 [14] with broad cell type markers examined to annotate cluster identities (Fig. 1B; Suppl. Fig. 1). Figure 1B presents a tSNE plot for the integrated PD, MSA, and Control samples showing integration of all samples and nuclei types with broad markers localized within each identified cell type cluster. For all putamen samples, oligodendrocytes were found to be the predominant cell type, and Fig. 1C shows boxplots with the proportions of nuclei of each type among samples, constituting (mean ± standard deviation (SD)): 70.3 ± 14.8%, 66.6 ± 25.3%, and 67.7 ± 13.5% of total sample nuclei, for Control, PD, and MSA samples, respectively. No significant differences in broad cell type proportion distributions were observed among conditions (X-squared p-value = 0.2759).

**Fig. 1.**
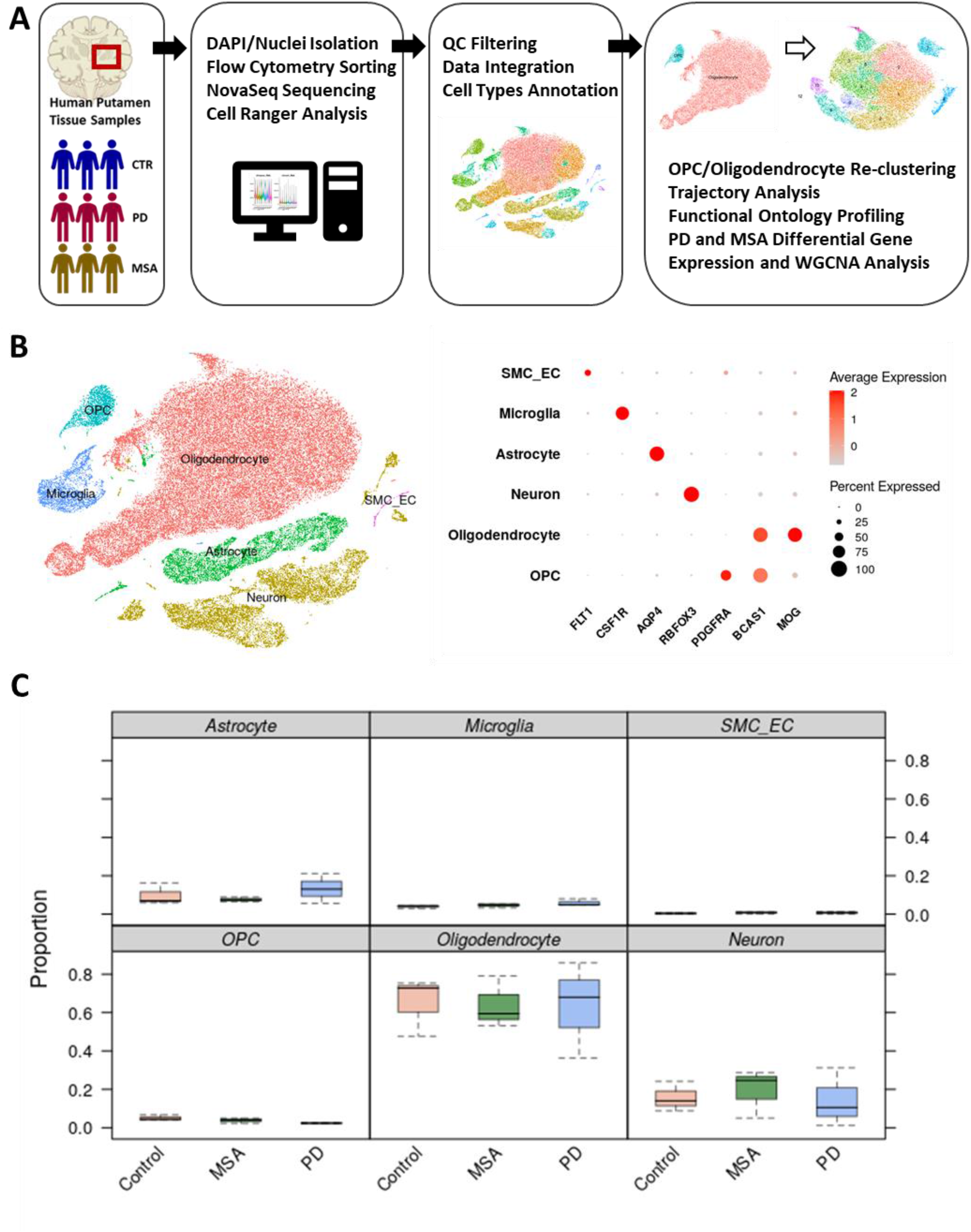
Oligodendrocytes are the major cell type in putamen. Schematic workflow for snRNA Seq analysis of human putamen samples (A). Data integration and clustered nuclei broad cell type marker expression identifies oligodendrocyte lineage as dominant type in putamen (B). Sample integration shown by condition and broad cell type nuclei proportions by sample. No significant difference exists in distribution of cell types by disease group (X-squared p = 0.2759) (C).

### Putamen Oligodendrocyte Populations Exhibit Molecular Diversity

Oligodendrocyte lineage nuclei were then subset from the integrated objects, to undergo re-clustering and profiling to identify cell subtypes, evaluate their relationships by pseudotime trajectory analysis, and identify differentially expressed genes between disease (PD or MSA) and Control nuclei. Among oligodendrocyte lineage nuclei, oligodendrocyte precursor cells (OPC; markers: PDGFRA/VCAN), committed oligodendrocyte precursor cells (COP), and multiple clusters of oligodendrocytes (MOG/MAG/OPALIN/HAPLN2) have been described in previous studies [4, 15]. Re-clustering of the OPC/Oligodendrocyte lineage cells was performed based on the rationale that unsupervised clustering of broad cell types might generate different cluster separations among oligodendrocyte subtypes than might be identified through applying the procedure only to oligodendrocyte lineage cells. Among 13 oligodendrocyte lineage clusters identified by unsupervised clustering, profiling of Control nuclei revealed similar maturation markers and pseudotime values for two COP clusters and five pairs of oligodendrocyte clusters (Supplementary Fig. 3). Clusters similar in their marker and pseudotime profiles were combined to yield seven clusters for downstream profiling: OPC, COP, OL1, OL2, OL3, OL4, OL5 (Fig. 2A; Suppl. Fig. 2).

**Fig. 2.**
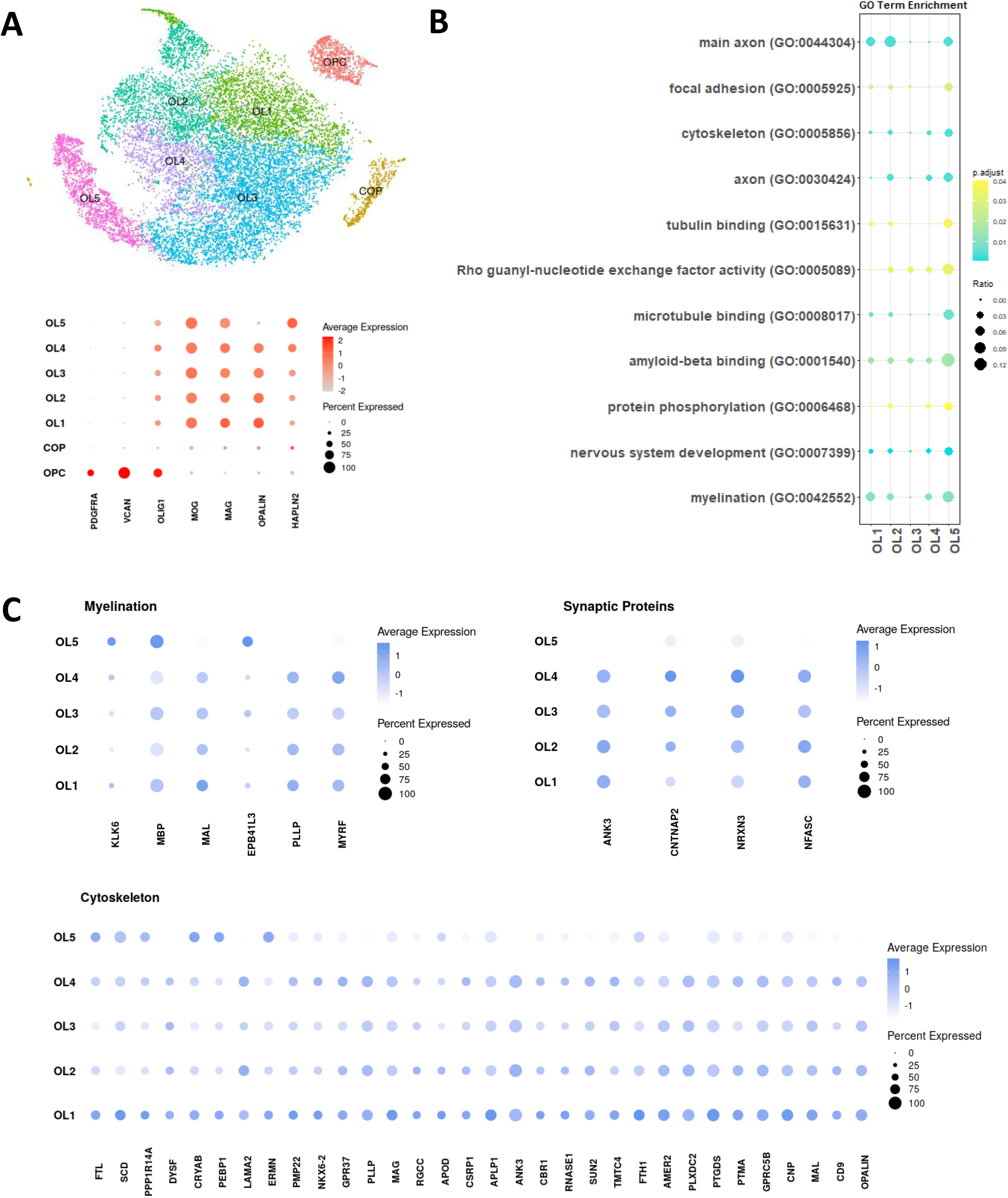
Oligodendrocytes subtypes in Putamen tissue. Clustered oligodendrocyte nuclei from CONTROL patients ordered by pseudotime and maturation marker expression (A). Differentiated oligodendrocyte subtypes, OLI 1-5 marker genes identified by MAST analysis (FC>1.2; adjusted p-value<0.01) are enriched for unique Gene Ontology Biological Process terms. Top enriched pathways are shown for adjusted p-value < 0.05. The ratio represents the proportion of term genes which are marker genes (B). Selected marker gene expression shown by cell subtype cluster for selected Gene ontology enriched terms (C).

Comparing labelled oligodendrocyte clusters OL1-5, marker genes for these clusters were identified using the MAST package with cutoffs fold change > 1.2, adjusted p-value < 0.05, and detected expression in 60% of cells. The list of all cluster marker genes was then then examined for Gene Ontology term enrichment to evaluate functional differences between these clusters. Selected highly significant term enrichments are shown in Figure 2B. Of note, OL1 was found to be enriched for genes involved in myelination (OPALIN, PLP1, MBP, MAL, EPB41L3, PLLP, MYRF, and NFASC), consistent with the results of trajectory and maturation marker analysis. Clusters with increased HAPLN2 expression were found to exhibit greater expression of gene transcripts involved in cytoskeleton regulation (DYNC11, DYNC12, KIF1A, KIF5C, ACTN2) and cell adhesion genes functionally important for neuronal guidance, formation of neuronal projection, and axonogenesis (e.g. STMN1, MAP1B, SEMA5A, EPHB2, S100B, PRKCA). Complete lists of oligodendrocyte cluster marker genes are included as supplementary files. Cluster markers also include a number of genes coding for synaptic proteins (NFASC, NRXN3, CNTNAP2, ANK3) which were most highly expressed in the middle trajectory oligodendrocyte clusters (Fig. 2C), in line with previous evidence of synaptic-like communication between neurons and oligodendrocyte lineage cells being important for myelination (Fig. 2C) [16–18]. Oligodendrocyte clusters were also notably enriched in small GTPase regulatory proteins which have been reported to play roles in regulating neuron projections and synapsis (RASGRF1, PLEKHG1, PREX2, EPS8) [19, 20]. Fig. 2C shows expression of marker genes for a subset of enriched functional pathways, illustrating cluster-level functional heterogeneity.

### Disease-Specific Alterations in Heterogeneity in the Putamen Oligodendrocyte Populations

After profiling Control oligodendrocyte functions by cluster, we then examined the relative distributions of PD and MSA nuclei among the integrated clusters (Fig. 3A) and performed within-cluster differential gene expression analysis using MAST [21]. Comparison of subtype proportions among conditions by X-squared testing revealed borderline significant differences in population distribution by condition (p = 0.05478; Fig. 3B). For all conditions, we observed the largest proportions of oligodendrocytes fell in the myelinating OL1 to mid-trajectory OL3 clusters, with lower proportions observed in the OL4 and OL5 clusters.

**Fig. 3.**
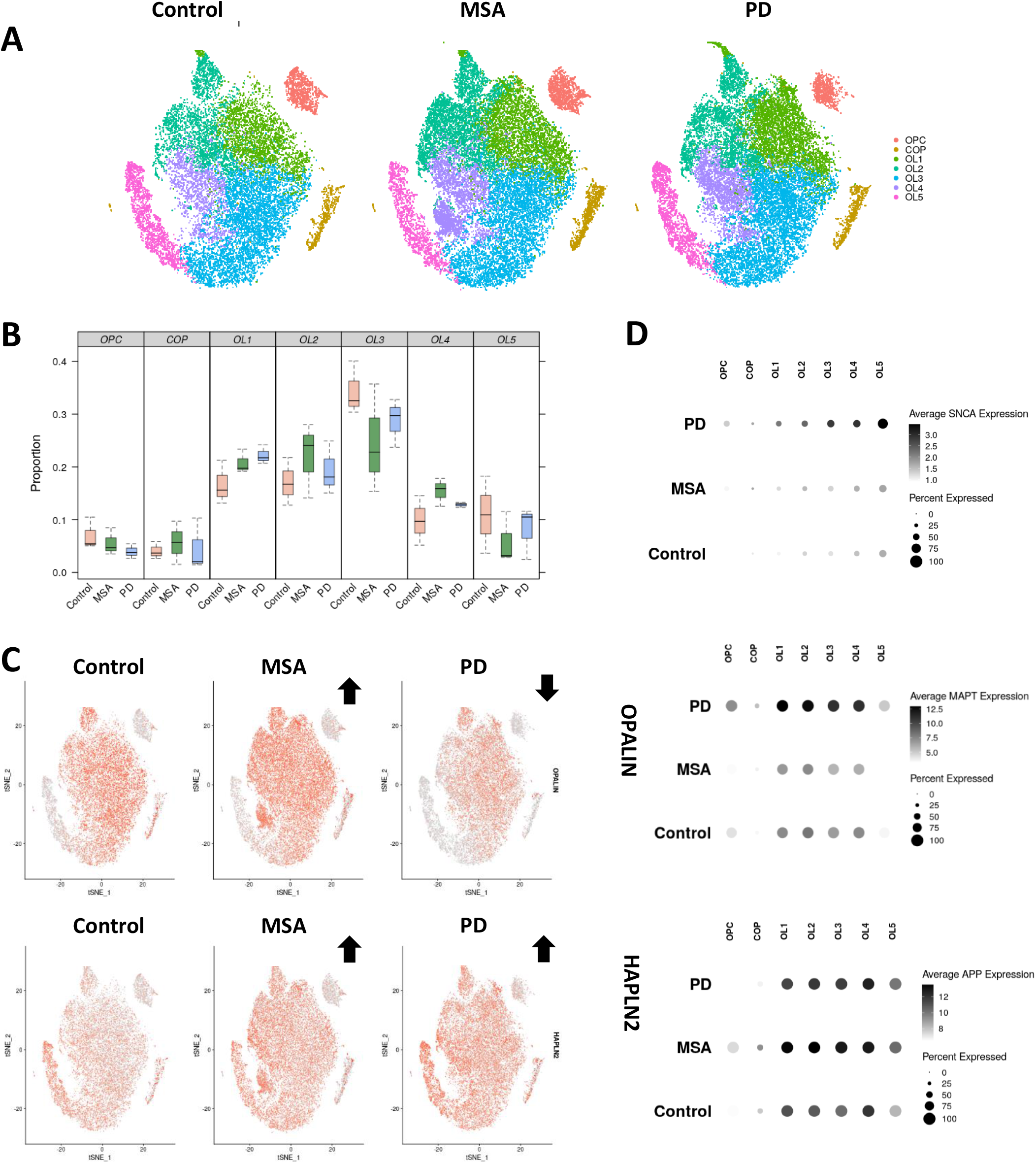
Analysis of oligodendrocyte heterogeneity in PD and MSA. Re-clustered oligodendrocyte lineage cells grouped into OPCs, COPs and five myelinating and mature OLs(A). Proportions of nuclei distributed among the clusters. Chi-squared test applied to oligodendrocyte proportions by cluster identifies a significant relationship between condition (Control, MSA, PD) and oligodendrocyte subtype proportions (X-squared=20.711, df=12, p-value = 0.05478) (B). Significantly decreased OPALIN and increased HAPLN2 expression is observed in PD versus Controls; OPALIN and HAPLN2 are increased MSA relative to Controls - low (grey) to high (red) expression (C). Neurodegeneration-associated genes APP, MAPT, and SNCA expression differs among oligodendrocyte subtypes (D).

Given the observed differences in cluster proportion distributions, we next proceeded to compare the expression of marker genes OPALIN and HAPLN2 among samples as this could provide some insight into whether oligodendrocytes in different conditions might be more actively transitioning between states. Interestingly, we found that PD oligodendrocytes exhibited significantly greater expression of HAPLN2 across multiple oligodendrocyte clusters in comparison to Controls and significantly reduced OPALIN expression (Fig. 3C, Suppl. Fig. 3). In MSA, however, both greater expression of OPALIN and HAPLN2 was observed compared with Controls (Fig. 3C, Suppl. Fig. 3). We then additionally examined whether expression of neurodegeneration-associated genes APP, MAPT, and SNCA between these oligodendrocyte subclusters (Fig. 3D, Suppl. Fig. 4), and we observed increasing average expression of SNCA in clusters with greater HAPLN2 expression. In PD, this increasing expression of SNCA was more pronounced than was observed for Control or MSA clusters. In addition, in PD, greater expression of MAPT was also observed, particularly in OL1-4 clusters, while APP expression was found to be increased in both PD and MSA (Fig. 3D, Suppl. Fig. 4).

**Fig. 4.**
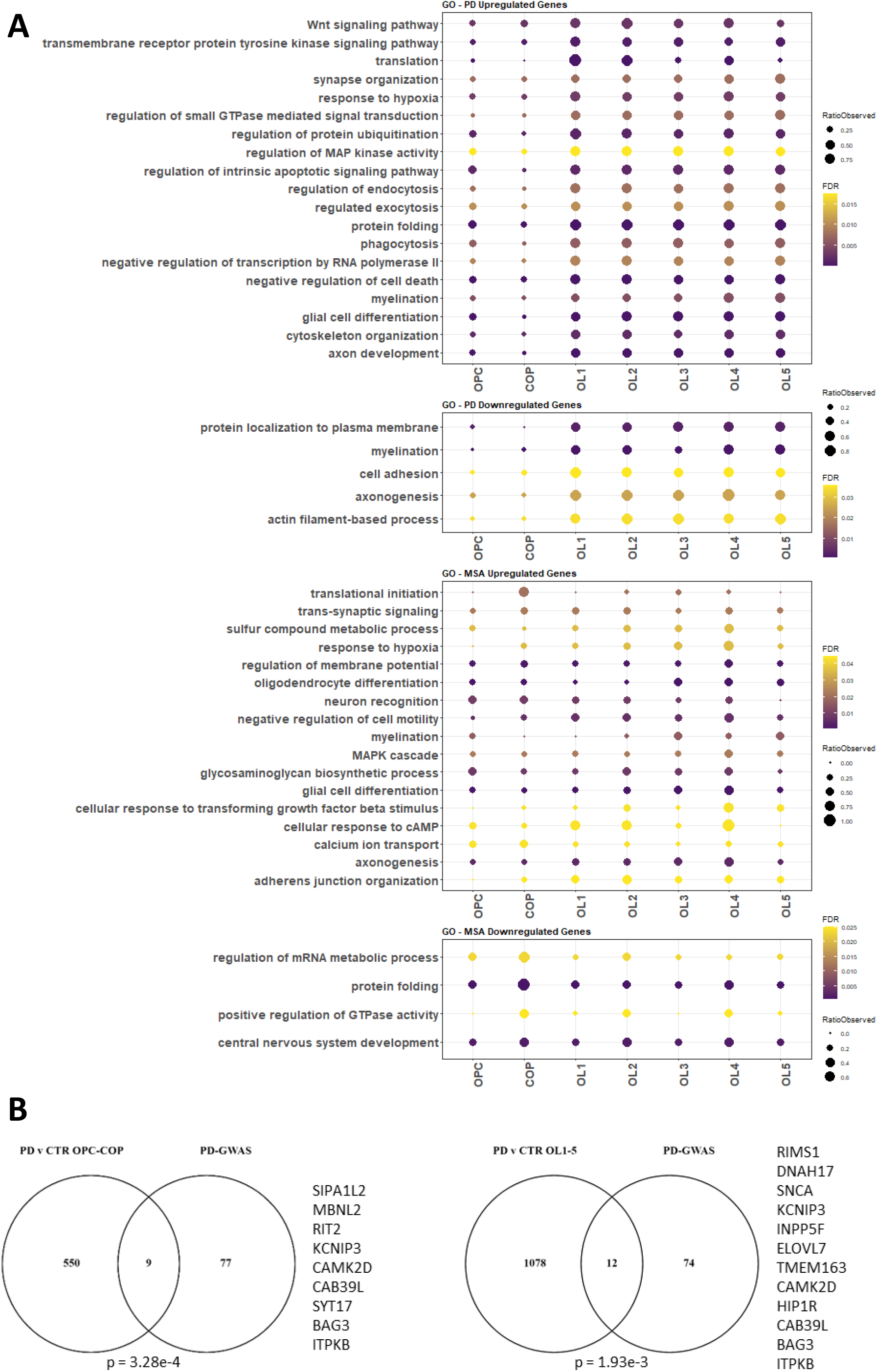
Differential gene expression analysis in PD and MSA oligodendrocyte subtypes. Differentially expressed genes compared by cluster for PD versus Control and MSA versus Control were analyzed for their associated Gene Ontology Biological Function terms (A). Overlaps among PD-GWAS nearest genes and differentially expressed genes in PD vs Control analysis(B).

### Distinctive PD- and MSA Transcriptomic Alterations in Putamen Oligodendrocytes

Differential gene expression analysis was then performed within each labelled cluster for comparisons of PD versus Control nuclei and MSA versus Control nuclei using MAST [21]. The list of differentially expressed genes as per cluster and disease condition is available as supplementary (Suppl. Fig. 5). Gene Ontology term enrichment analysis was performed using STRING [22] for pairwise comparison using a cut-off adjusted p-value at least < 0.05. Figure 4A shows GO term enrichment z-scores for selected categories for upregulated and downregulated gene sets compared for PD versus Control and MSA versus Control. Comparison analysis by cluster revealed both common and distinctive impacted pathways for PD and MSA compared with Control (Fig. 4A; Suppl. Table 3; Suppl. Fig. 5). We observed in PD, a general upregulation of genes responsible for protein folding and disaggregation across the groups. These include several cytosolic and ER resident members of the HSP70, HSP90, HS40, CCT and FKBP protein chaperon families. Interestingly, many of these proteins are well known targets of HSF1 or ATF6 transcription factor [23, 24]. Also upregulated in several oligodendrocyte subtypes in PD are transcripts encoding for multiple ribosomal proteins, tyrosine kinase (IGF1R, KIT, PRKD1, NRG2, NRG3, IRS2, GRB10, SOS1, FGF1) and small GTPase (ARHGAP17, ARHGAP32, RHOBTB1, ARHGAP12, RAC1, ARHGEF7, ARHGAP5, ARHGAP35) signaling regulators a well as proteins involved in vesicular transport (SNX3, MIB1, MFGE8, HIP1, SNCA, CAV1, RAC1, RAB14, LDLRAP1, PICALM). Among genes downregulated across lineages in PD were genes connected to neurogenesis, including several genes involve in axonogenesis (SEMA6A, ANK3, PTK2, AUTS2, FGFR2, ROBO2, ROBO1, SLIT2, ZEB2, SEMA3B) and myelination (PLLP, JAM3, MAL, EPB41L3, CD9, MAG, MBP); phagocytosis pathway genes were observed to be highly expressed in PD but not MSA or Control nuclei (Suppl. Fig. 5).

**Fig. 5.**
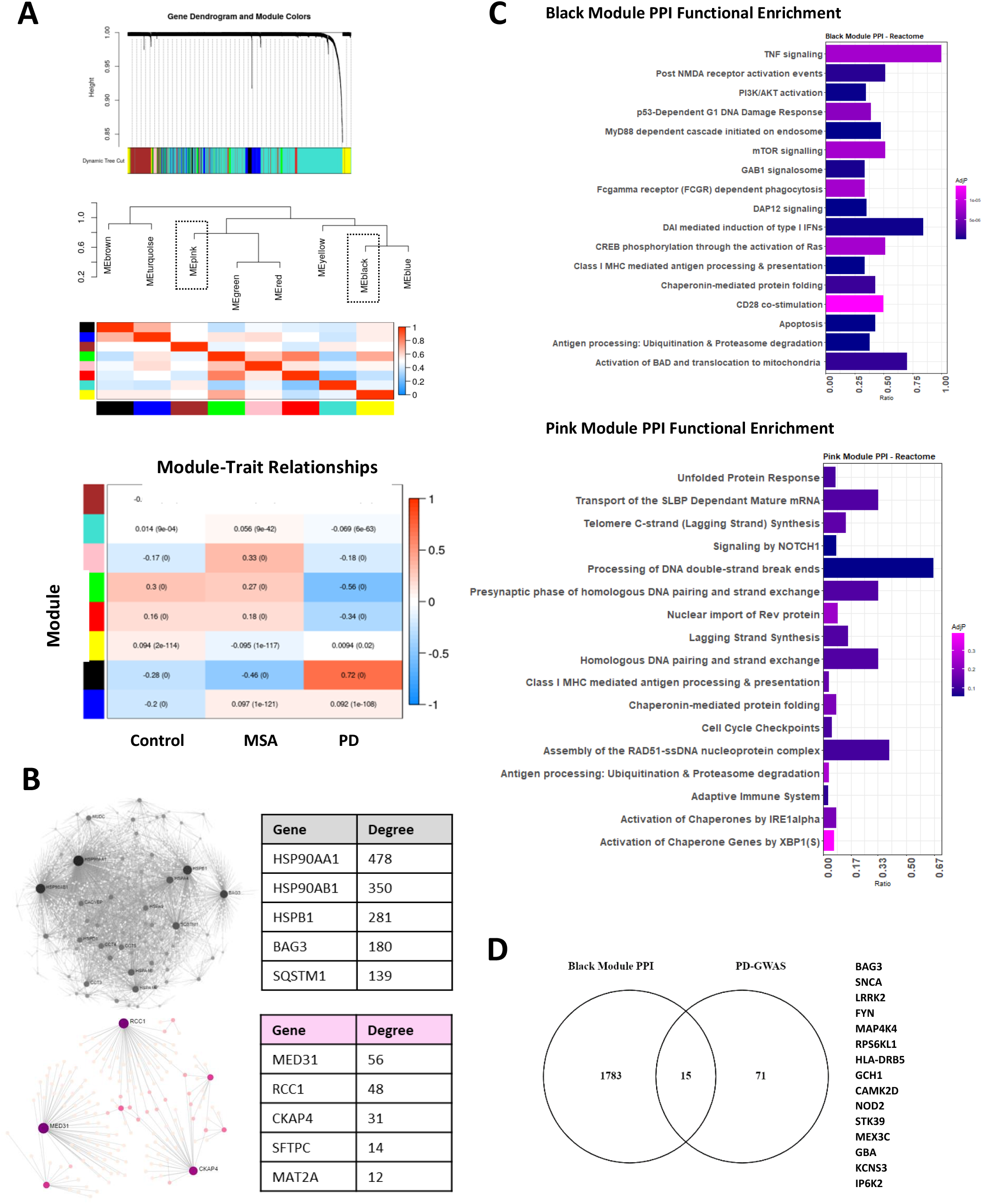
WGCNA. identifies co-expression modules significantly associated (|corr coefficient| > 0.3 for any of the comparisons and p-value <0.001) with PD (black, red, green) and MSA (pink) (A). PPI network build with indicated module gene as seeds. Selected network hub genes are shown (B). PD-associated black module and MSA-associated pink module PPI network members enrichment for GO functional terms(B). PPI network hub genes are shown (C). Black module PPI is significantly enriched for genes nearest PD-associated GWAS variants nearest genes (p = 0.01046) (D).

In notable contrast to PD, fewer pathways were found to be dysregulated in MSA oligodendrocytes when compared to Controls. Multiple MSA oligodendrocyte lineage clusters exhibited significantly reduced expression of genes such as HSPA1A, HSPA1B, DNAJB1 and FSKBP5 which are involved in protein folding (as opposed to PD where gene expression within this functional category is upregulated). Within the OPC cluster, we also observed the downregulation of genes involved in regulation of apoptosis and senescence (EGFR, NFKB, STAT3, FOXO1, MDM2, CDKN2A, TNIK, TXNIP and BCL6). In more mature oligodendrocytes, we observed increased expression of several proteins associated with neurogenesis, including several involve in axonogenesis (ROBO2, SLIT2, EPHA3, THY1, LRRC4C, NRNX1, CTNNA2, DPYSL2, NRP2), synaptogenesis (GRIA2, GRIA4, GRIN2A, NLGN1, DLGAP1, LRRTM3, LRRC4C, LRRTM4, LRRC7, VAMP1, GABRB1, GRIK2) and oligodendrocyte differentiation and myelination (SOX6, NKX6-2, OLIG1, KCNJ10, CNP, OMG, CD9, PARD3 PNP22, MPZ, MOG, ERMN and THRB). (Fig. 5, Suppl. Table 3).

We then compared differentially expressed genes in oligodendrocytes with genes linked with highly significant variants identified by GWAS for PD [25] and genes linked with recent potentially significant variants in a recent GWAS for MSA [26]. For PD, we observed significant enrichment for PD-GWAS genes among differentially expressed genes for both OPC-COP (hypergeometric p = 3.28e-4) and OL1-5 (hypergeometric p = 1.93e-3) (Fig. 5B). No enrichment was observed among differentially expressed genes linked with intermediate significance variants for the reference MSA GWAS [26].

### Identification of Disease-Correlated Gene Modules by WGCNA

In addition to the within-cluster analyses of differentially expressed genes, we also used WGCNA to examine whether there might be modules of coregulated genes associated with each disease, independent of subpopulation. WGCNA was applied to the nuclei-count matrices for the integrated PD-MSA-Control object, with the identification of modules of co-expressed genes specifically correlated with PD and with MSA which are labelled by color (Fig. 5A). Several modules were strongly and significantly correlated with disease condition. The Black module positively correlated with PD (corr 0.72; p<1e-5) and negatively correlated with Control and MSA, and the Pink module positively correlated with MSA (corr 0.33; p<1e-5) but showed a weak but significant negative correlation with both Control and PD. Modules Green and Red showed a strong negative correlation with PD and a positive correlation with both Control and MSA (Fig. 6A). In line with the above-described differential expression analysis (Fig. 5), a significant fraction of the black module 103 genes have functions related to the cellular response to stress, including genes involved in protein folding and the regulation of cell death. The modules Green and Red contain genes related to myelination, axonogenesis, synapsis, alternative splicing, and regulation of the cytoskeleton (Suppl. Fig. 6). The Pink module only contained 38 genes, suggesting greater transcriptomic profile similarities for MSA in comparison with the Control group.

**Fig. 6.**
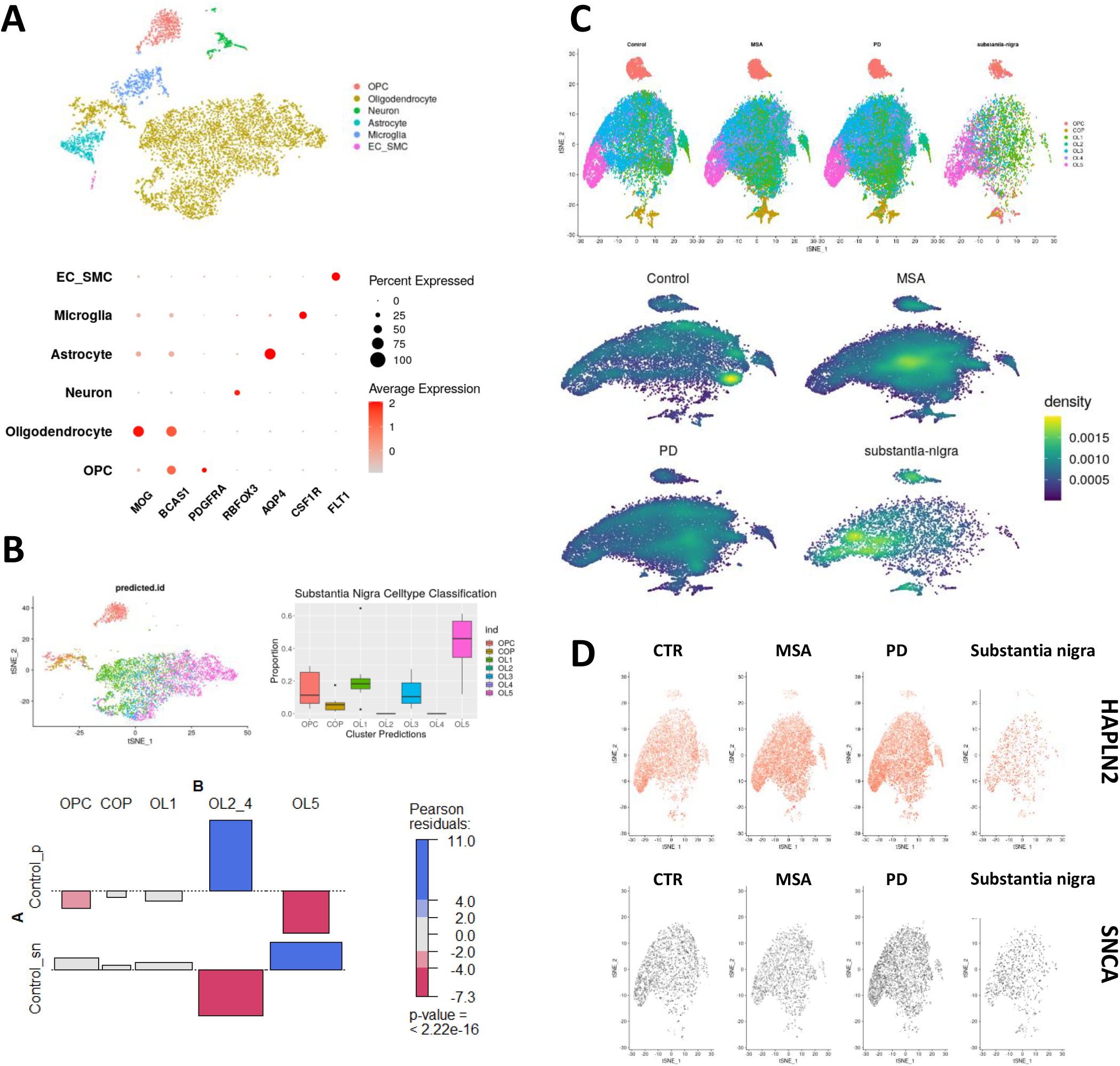
Broad cell type annotation and marker expression level for substantia nigra samples. From donors without neurological disease (n = 5 donors) (A). Label transfer to substantia nigra oligodendrocytes from putamen predicts that substantia nigra contains significantly higher proportions of OL5 type oligodendrocytes and fewer mid-trajectory oligodendrocytes compared with Control samples (p<2.2e-16) (B). Reintegration of all samples confirms greater density of OL5 oligodendrocytes among substantia nigra nuclei(C). SNCA expression in oligodendrocytes of substantia nigra is not observed to be higher in regions expressing oligodendrocyte maturation marker HAPLN2 (in contrast to PD) (D).

To better understand the possible functional consequences of the transcriptional changes associated to the disease-related modules of coregulated genes, we generated tissue-specific protein-protein interaction networks for genes included in the Black, Red, Green and Pink modules. Top hub genes for the module-seeded PPI networks for the Black and Pink modules are presented in Fig. 6B and Suppl. Fig 6 (Red and Green modules). Complete module gene lists and PPI network files are included as supplementary files, labelled by color. Pathway enrichment profiling for each PPI network was then performed using Reactome [27]. Consistent with differential expression analysis, enriched pathways in the PD-associated Black module PPI network most prominently included genes involved in protein folding, stress, and regulated apoptosis. In contrast, the Pink module, which was positively associated with MSA, was more strongly enriched for functions related DNA replication, cell cycle regulations, and DNA damage repair (Fig. 6C). The Green and Red modules (negatively correlated with PD and positively correlated with MSA and Control) were notably enriched for functions including chromosome maintenance and DNA damage repair functions (Green) as well as gap junction trafficking (Red) (Suppl. Fig. 6).

We also investigated whether genes in the Black, Pink, Red, and Green module-associated PPI networks might be causally linked with PD pathogenesis. To do this, we compared genes in these PPI networks with a list of nearest genes to highly significant variants linked with PD risk by GWAS, as we had done for cluster differentially expressed genes [25]. Genes in module-associated PPI networks linked to disease risk by genetic evidence are particularly interesting genes as these may contribute to disease-specific pathobiological mechanisms. Statistical testing for overlap enrichment identified significant Black module PPI enrichment for nearest genes to highly significant SNPs linked with PD by GWAS (15 genes; enrichment test p-value = 0.01046). (Fig. 6D). We also compared genes in the Pink, Red, and Green module PPI networks with a list of genes linked with intermediate significance variants in a recent MSA GWAS [26]. Among these, the single gene from this MSA-GWAS list found in the Red, Green, and Pink PPI networks was for ELOVL7, which was identified in the Green module PPI network.

### OL5 Oligodendrocytes Are the Major Type in Substantia Nigra

As substantia nigra is specifically impacted in PD and involved to varying degrees in MSA, we then integrated putamen oligodendrocyte nuclei from Control, MSA, and PD samples with an independent external snRNA-seq data set for substantia nigra samples obtained from individuals without neurological disease [3]. This integration was performed to more directly relate our findings to oligodendrocyte functional biology in this target region. After sample integration and broad cell types annotation for substantia nigra samples (Fig. 6A; Suppl. Fig. 8), oligodendrocyte and OPC nuclei were separated. We then separately applied two procedures to integrate substantia nigra samples with profiled putamen data: label transfer and re-integration. By label transfer, 43.1 ± 17.7% of substantia nigra nuclei were classified as OL5, a significantly different distribution on comparison with Control putamen samples (X-squared p-value<2.2e-16) (Fig. 6B). To secondarily confirm this classification, we re-integrated all putamen and substantia nigra samples and examined whether these predicted labels aligned in this new integration (Fig.6C). Density plots confirmed differential concentration of substantia nigra oligodendrocytes near OL5 oligodendrocyte clusters following CCA-based integration (Fig. 6C). As noted above, this OL5 oligodendrocyte cluster is the one with the highest HAPLN2 expression in putamen samples and the cluster with the greatest overexpression of SNCA in PD. SNCA expression in oligodendrocytes of healthy substantia nigra was in regions expressing marker HAPLN2 was observed to be most similar to Control putamen (in contrast to PD) (Fig. 6D).

## Discussion

The role of oligodendrocytes in neurodegeneration has received less attention in the past compared to their study in relation to the classical demyelinating diseases. In this study, we performed snRNA-seq profiling of oligodendrocyte populations in human putamen samples from Control, PD, and MSA donors. We found that oligodendrocytes were the dominant cell population for all samples, and we profiled 7 subpopulations identified by trajectory and differential expression analyses (Fig. 1,2). Using Control nuclei as a reference, we identified functional enrichment patterns for Control oligodendrocyte subpopulations, including ontology enrichment for myelination in OL1, expression of synaptic protein genes in OL1-4, and vesicle and cytoskeleton-mediated transport activities in OL5, among others (Fig. 2). While myelination is the most well-recognized function of oligodendrocyte lineage cells, activities including synaptic signaling, metabolic support, and immune and inflammation pathway modulation by OPCs and oligodendrocytes are increasingly being recognized [28–30], and subpopulations enriched for these activities are represented among our results.

Recent developments in single cell and single nuclei sequencing have paved a way for more comprehensive studies and profiling of oligodendrocytes in different brain diseases [3, 7, 15]. A novel finding in this study is our identification of a subpopulation of oligodendrocytes expressing greater levels of HAPLN2 (OL5), which we identify as the predominant oligodendrocyte subtype in healthy substantia nigra (Fig. 6) and which is also found to be the cell population with the greatest level of SNCA expression in the putamen in PD. Hyaluronan and proteoglycan link protein 2 (Hapln2) also known as Brain-derived link protein 1 (Bra1) is expressed in myelinated fiber tracts in the adult brain, where this protein contributes to the binding of proteoglycans to hyaluronan in the extracellular matrix, stabilizing the extracellular matrix, providing diffusion barrier support particularly around the nodes of Ranvier, and maintaining signal conductivity [31]. Deficiency of Hapln2 alters extracellular matrix protein expression and results in nerve conduction disturbances [32]. HAPLN2 overexpression in substantia nigra has been previously reported as a feature of PD [32, 33], and we observe such overexpression in PD as well as MSA oligodendrocyte populations. Significantly, HAPLN2 overexpression has also been shown to result in α-synuclein aggregation and neurodegeneration in an experimental animal model of PD [34]. While we observe of greater numbers of HAPLN2-expressing oligodendrocyte types in substantia nigra and greater expression of HAPLN2 in both PD and MSA oligodendrocytes, it remains to be further understood how alpha-synuclein might be interacting with Hapln2 in this region and well as what might stimulate HAPLN2 expression in each of these disorders.

SNCA gene expression was found to be significantly higher in PD oligodendrocytes versus Control as well as in comparison to MSA (Fig. 3D). While oligodendroglial cytoplasmic inclusions of α-synuclein protein are described as the predominant neuropathological finding in MSA and neuronal α-synuclein aggregates are described as being more prominent in PD, varying degrees of neuronal and oligodendroglial involvement are reported in both disorders [35–38]. SNCA mutations, duplications, and triplications have been causally linked with familial PD in multiple studies [39–41]. While genetic variants within the SNCA locus have also been associated with MSA in a few studies [42, 43], the connection between SNCA overexpression in oligodendrocytes and MSA is less clear. Cell-to-cell transmission of highly pathogenic misfolded α-synuclein proteins from neurons to oligodendrocytes has been posited as one potential explanation for the prominent oligodendroglial inclusions observed in MSA [44], and our observation of lower levels of oligodendrocyte SNCA expression in MSA versus PD may lend some further support to this theory [31–34, 45].

Differential gene expression analysis and WGCNA provide further insight into differences in oligodendrocyte involvement in PD versus MSA. We observed both shared and distinctive transcriptional alterations (in PD versus MSA Oligodendrocytes), including more prominent upregulation of gene transcripts related to cellular response to stress in PD versus MSA. Specifically, several cytosolic, mitochondrial and ER resident chaperone and co-chaperone proteins, modulators of JNK stress kinase activation, several tyrosine kinase receptors and apoptosis regulators were upregulated in PD as compared to MSA, where the same genes were either downregulated or did not change expression in MSA versus Controls. Given the importance of cellular protein folding machinery to counteract proteotoxic stress created by protein inclusion bodies and aggregates such as those associated with synucleinopathies, this downregulation in MSA may contribute to disease pathology and may help to explain why SNCA accumulates in MSA but not in PD oligodendrocytes. Heat shock proteins are both sensors and effectors of the cellular response to proteotoxic stress. In normal conditions Hsp90 and HSP/C70 proteins in the cytosol and HSP5A (also known as GRP78) respectively bind and inhibit the activity of HSF1 and ATF6. Accumulation of misfolded proteins releases this inhibition, leading to the activation of these transcription factors which in turn increases the transcription of heat shock proteins to alleviate stress by contributing to de novo protein folding, disaggregation and degradation of misfolded proteins. Prion-like propagation of endogenous α-synuclein species via axonal projections may explain activation of the heat shock response in putamen oligodendrocytes but also microglia in PD [2]. There are several possibilities as to why oligodendroglia in MSA putamen fail to display a proteotoxic stress response even though these cells can accumulate SNCA-containing cytoplasmic inclusions. Cytoplasmic glial inclusions are composed of hundreds of proteins including entrapped heat shock proteins [46], whose entanglement may help to explain the disruption of the heat shock response. Posttranscriptional mechanisms of negative regulation of transcription factors HSF1 and ATF6 could also hamper the response in MSA. Finally, high order aggregation of SNCA into inclusion bodies such as those found in MSA oligodendroglial cells may be a protective mechanism against the cytotoxic effects of SNCA fibrils [47], and the absence of a classical proteotoxic stress response in MSA oligodendrocytes may reflect this alternative mechanism of stress reduction. Understanding these mechanisms may have translational implications because several HSF1 pharmacological activators are being tested in clinical trials [48] that could be repurposed for MSA treatment. Noticeably, several other genes regulating other aspects of the stress response such as the autophagy receptor Sequestosome 1, redox enzyme SOD1, JNK1 kinase activator MAP2K4, chromatin condensing factor HP1BP3k, and Sirtuin 2 follow a pattern of expression like that of the Heat Shock proteins. Given the complexity of the proteotoxic stress response in MSA and PD, additional studies on how these changes affect OPC differentiation, myelination and oligodendrocyte survival are warranted.

Immunohistochemical analysis of PDGFRa+ cells have demonstrated an elevated number of OPCs in the striatum of MSA patients [49]. Because MSA is accompanied by myelin pallor and reduced expression of MBP [50], a defect in oligodendrocyte maturation has been proposed as a feature of this disease. Our results serve to qualify this hypothesis. Firstly, we observed no increase in the number of OPCs in MSA or PD putamen but in the case of MSA, these OPCs exhibited a modest but significant increased expression of OPC marker PDGFRa and oligodendroglia differentiation factors, OLIG1, OLIG2 and MYT1. No significant differences in the number of myelinating and mature oligodendrocytes are obvious in MSA and, except for a slight reduction in MBP expression in COP, no diminished expression of myelin proteins was obvious. Instead, increased expression of genes involved in glial differentiation and myelination (NKX6-2, OLIG1, KCNJ10, CNP, OMG, CD9, PARD3 PNP22, MPZ, MOG, ERMN and THRB) is observed in myelinating and mature oligodendrocytes. Interestingly, SOX6 a transcription factor that can inhibit the expression of myelin genes when overexpressed in mice is upregulated in these cells [51]. In contrast, PD oligodendrocyte expression of several genes involve in myelination, including OPALIN, MBP, MOG, PLLP, MAL, MAG, MOBP, as well as involved in cell-cell adhesion, cell projection organization and in lipid synthesis (SREBP1, FADS1,2, ELOV6 ad INSIG1), functions important for efficient myelination, are downregulated in PD oligodendrocytes. Reduced expression of these genes suggests diminished potential for myelination activity. White matter loss is an early feature of several neurodegenerative disorders including Alzheimer’s Disease and vascular dementia [52] and has been increasingly identified among PD patients [53]. These results further suggest that reduced myelination activity may also occur in PD.

When the genes dysregulated in PD putamen were analyzed for the presence of SNPs associated with increased risk of PD in GWAS studies, we found 12 genes (RIMS1, DNAH17, SNCA, KCNIP3, INPP5F, ELOVL7, TMEM163, CAMK2D, HIP1R, CAB39L, BAG3, ITPKB) containing such polymorphisms. These genes include SNCA as the only well-recognized PD gene regulated in OL. We also observed dysregulation of MAPT, a recognized genetic risk factor for PD [54]. Our search of genes linked with intermediate significance variants from a recent MSA GWAS [26] found only one gene, (ELOVL7). ELOVL7 is a transferase which catalyzes long chain fatty acid synthesis [55]. ELOVL7 coding variants are rare, and not all studies have identified a consistent association of this gene with MSA. Transcriptomics analyses, although comprehensive can only provide a limited view of the regulatory mechanisms in volved in PD and MSA etiology. A significant amount of such mechanisms occur postranslationally and cannot be easily predicted from mRNA level changes alone. We use network analysis to study the possible repercussions of the transcriptional changes we observed. Protein functional interaction networks capture the functional relationships (edges) between nodes (proteins) through integration of multiple datasources. Therefore, we built funtional protein interaction networks using as seeds the sets of coexpressed genes (modules) that covariated with disease status. PPI network generated for the PD-associated Black module showed significant enrichment for genes linked with highly significant variants identified by GWAS analyses for PD [25]. including several genes strongly and significantly associated with PD risk, including GBA, STK39, LRRK2, and others. In contrast, modules positively correlated with MSA were not significantly enriched for genes linked with intermediate significance variants from a recent MSA GWAS [26].

In summary, we provide a detailed analysis of transcriptional changes at a single cell resolution in Putamen oligodendroglia for PD and MSA patients. Distinct transcriptional changes suggest that PD and MSA, while both synucleinopathies, possess important pathophysiological differences in machineries for cellular response to stress and for myelination activity. Future work, including the analysis of greater number of patient samples is needed to verify and generalize our observations, and further studies are also needed to examine how gene expression changes relate to protein levels by orthogonal analytic methods. The oligodendrocyte subpopulations profiled here may exhibit distinctive functional activities which may offer promising therapeutic targets for these debilitating and often lethal diseases.

## Methods

Fig. 1A presents the analysis workflow. Post-mortem fresh-frozen unfixed human putamen samples were each obtained through partnerships with licensed organizations with completed pre-mortem consent for donation and ethical committee approval for sample acquisition and use. Samples used for single-nucleus RNA sequencing (snRNA-seq) were putamen tissue sections from nine human donors (n =3 per group, mean age in years ± SD: Control, 78.7 ± 9.5; PD, 79.7 ± 5.5; MSA, 65.0 ± 10.6) (Supplementary Table 1).

### Nuclei Isolation

Tissue samples were stored at −80°C. For tissue lysis and washing of nuclei, sample sections were added to 1 mL lysis buffer (Nuclei PURE lysis buffer, Sigma) and thawed on ice. Samples were then Dounce homogenized with PestleAx20 and PestleBx20 before transfer to a new tube, with the addition of additional lysis buffer. Following incubation on ice for 15 minutes, samples were then filtered using a 30 mM MACS strainer (MACS strainer, Fisher Scientific), centrifuged at 500xg for 5 minutes at 4°C using a swinging bucket rotor (Sorvall Legend RT, Thermo Fisher), and then pellets were washed with an additional 1 mL cold lysis buffer and incubated on ice for an additional 5 minutes. Samples were then centrifugated at 500g for 5 minutes at 4°C and then were resuspended in 1mL Nuclei PURE Storage Buffer (Nuclei PURE storage buffer, Sigma). Sample washing was performed until the supernatant cleared. A final resuspension was then prepared in 0.6mL wash buffer before NeuN/Dapi staining and FACS sorting was performed. For NeuN/Dapi and FACS sorting, from 0.6 mL nuclei sample, 540 mL, 30 mL, and 30 mL were aliquoted into tubes for sample and controls and then 10X Dapi/NeuN buffer was added to tubes for a final 1X concentration. Tubes were then incubated on ice for 30 minutes, with inversion every 10 min. Following incubation, samples were spun at 500xg for 5 min, supernatant removed, and samples were resuspended in 600 ul Wash buffer for samples (300 ul for control tubes). Nuclei then underwent filtering and sorting using BD Bioscience InFlux Cell Sorter.

### Library preparation and NovaSeq Sequencing

Libraries were prepared according to 10xGenomics protocol for Chromium Single Cell 3’ Gene Expression V3 kit. NovaSeq sequencing was performed according to illumine NovaSeq 6000 protocol. UMI count matrices generated by Cellranger V3.0.2.

### Putamen Data Preprocessing

Summary information for final UMI count matrices for nuclei by individual sample and sequencing data are presented in Supplementary Table 2. Count matrices together with nucleus barcodes and gene labels were loaded with R version 3.6.1/RStudio for sample integration and unsupervised clustering using Seurat Package version 3.1. For Quality Control (QC), nuclei were filtered following standard protocols based on examination of violin plots. Cutoffs 200 < nFeature_RNA < 9000 and percent.mt < 5 were used. Filtered matrices were then individually log-normalized by sample according to standard Seurat workflows. After quality filtering, 87,085 total nuclei were included in the final dataset. Individual-level post-QC nuclei and feature counts are presented by sample in Supplementary Table 2 and QC filtering plots in Supplementary Fig. 1.

### Broad Cell Types Annotation

Sample integration was performed in Seurat using the FindIntegrationAnchors and IntegrateData functions for 2000 variable features. Following integration and scaling according to Seurat package workflows, a range of clustering resolution values were trialed prior to broad cell type annotation (trial parameters: 2000 variable features, nPC = 20, and resolution = [0.6, 0.8, 1.2]; TSNE plots for different trial parameters included as Supplementary Figure 2). Cluster-level expression of major cell type markers was examined and used to annotate cells contained within each cluster. Clustering resolution 0.8 was selected as our final clustering parameter. This value was found empirically to achieve optimal separation and consistency of cell type marker expression.

### Oligodendrocyte Lineage Re-Clustering and Profiling of Control Oligodendrocyte Lineage Nuclei

Nuclei identified as belonging to oligodendrocyte lineages by canonical marker expression were subsetted from the Seurat object by cluster name (“OPC” and “Oligodendrocyte”) (Control, n = 15,629 nuclei; PD, n = 20,839 nuclei; MSA, n = 21,799 nuclei) and this subset Seurat object was re-clustered using a range of clustering parameters: nfeatures = 1000 and resolution = [0.4, 0.6, 1.0]. For oligodendrocyte lineage re-clustering, nPCs = 15 and resolution 0.6 was chosen, as this value was found empirically to generate multiple clusters expressing different lineage markers (Supplementary Figure 1). All clusters generated contained nuclei from Control, PD, and MSA samples.

To profile Control oligodendrocyte lineage subpopulations for benchmark comparison with PD and MSA samples, Control nuclei were subsetted from the integrated Seurat object. Cluster expression of oligodendrocyte maturation markers was performed and compared with results from Slingshot trajectory analysis to order nuclei by developmental stage [56, 57]. Initial clustering identified 12 clusters. Several clusters classified as COP-type and multiple clusters of mid-trajectory oligodendrocyte clusters were found to have both consistent maturation marker profiles and similar slingshot pseudotime coordinates; these highly similar clusters were combined into a single COP cluster comprising clusters COP1-4 and middle trajectory oligodendrocyte clusters with consistent marker expression patterns were combined into cluster OL2 (Supplementary Figure 2). These cluster labels were applied to both the composite Seurat object including PD, MSA, and Control samples as well as the Control-only object.

### Differential Gene Expression and Pathway Enrichment Analysis

Genes differentially expressed in by each oligodendrocyte lineage clusters may relate not only to differentiation stage, but also potential functional and metabolic specifications. To profile cluster-level pathway enrichment patterns, cluster markers for oligodendrocyte nuclei were identified using the Seurat FindMarkers function with the MAST package [21]. We excluded OPC and COP clusters from this marker analysis as the distinctions of these clusters are previously characterized in published literature and including these nuclei in marker analysis might potentially conceal more subtle expression differences among differentiated oligodendrocyte subgroups. Markers used for downstream functional ontology analysis were those with a minimum percent expression 60%, adjusted p-value < 0.05, and fold change 1.2. Functional ontology analysis by cluster was performed for each cluster marker set using Enrichr (enrichment adjusted p-value < 0.05) [58].

Comparison of oligodendrocyte population distributions for PD, MSA, and Control samples was then performed. Proportions of nuclei in each group were calculated as the proportion of the total group nuclei. Differences in gene expression of the oligodendrocyte maturation markers HAPLN2 and OPALIN and neurodegeneration-associated genes SNCA, APP, and MAPT were assessed using the Seurat package visualization functions FeaturePlot() and DotPlot(), splitting nuclei by condition. Differentially expressed genes for PD versus Control and MSA versus Control were identified within each cluster using the Seurat FindMarkers() function and the MAST package for each pairwise comparison. GO term enrichment analysis among all differentially expressed genes was performed for upregulated and downregulated genes in comparison to Control for both PD and MSA.

### Weighted Gene Co-expression Network Analysis (WGCNA)

WGCNA was performed in R using the WGCNA package [59]. WGCNA was applied to the gene-gene co-expression matrix generated from the normalized cell-gene arrays from the integrated Seurat object. For gene co-expression modules identified by WGCNA, we calculated Pearson correlations between module eigenvectors and dummy-encoded disease state (PD, MSA, or Control) in order to assess if any of these modules might be significantly correlated with any particular disease state(s).

Positively correlated modules are those for which module genes are co-expressed more actively among nuclei from the correlated group, while negatively correlated module genes have relatively suppressed co-expression patterns. Module co-expression patterns may represent disease processes or possible cellular responses to synucleinopathy progression. To analyze disease-associated modules, we generated protein-protein interaction (PPI) networks seeded with module genes and performed pathway enrichment analyses using NetworkAnalyst and the STRING database with GO BP to generate and visualize PPI networks [60]. Network functional enrichment was queried using GO Biological Process terms.

PPI network genes were also compared with nearest genes to highly significant variants reported by Nalls and et al. in their 2019 meta-analysis of genome wide association studies for PD (PD GWAS). [25]. Statistical tests for overlap enrichment were performed in R using the hypergeometric test. Overlap visualizations for each of these steps were generated using Venny [61].

### Integration with Independent Substantia Nigra Data

Data was downloaded from supplementary files from Agarwal et al. 2020 for substantia nigra samples (5943 nuclei) obtained from five human donors and sequenced using the 10x Genomics Chromium Platform [3]. These files are accessible online through the NCBI interface: https://www.ncbi.nlm.nih.gov/geo/query/acc.cgi?acc=GSE140231.

Single-nucleus RNA-Seq integration and broad cell type annotations were performed for Substantia Nigra data separately in R using the Seurat package, version 3.0. A similar workflow as for putamen samples was followed. Pre-processing cut-offs were selected based on initial QC plots: 200< nFeatureRNA < 6000 and percent.mt < 5. Data were normalized at the individual sample level and then integrated using the Seurat functions FindIntegrationAnchors and IntegrateData as described in the Seurat data integration workflow with the number of PCs used for clustering (n = 20) chosen to optimize separation between clusters. Broad cell types were assigned for each cluster based on marker expression levels as for putamen. Clusters identified as OPC or Oligodendrocyte were subsetted from the combined object.

Label transfer was performed in Seurat using the FindTransferAnchors and TransferData functions to predict substantia nigra nuclei type. As a second validation step for these classifications, we also separately reintegrated all samples in Seurat and examined label and density distributions in tsne projects. Putamen and substantia nigra data oligodendrocyte lineage nuclei were integrated at the individual sample level with putamen sample data using the Seurat functions FindIntegrationAnchors and IntegrateData as described in the Seurat data integration workflow. For visualization, OPC and Oligodendrocyte nuclei from substantia nigra were labelled using their predicted cell type labels. Putamen sample nuclei retained their labels from earlier cluster profiling as described in the Methods section above. Samples were not re-clustered but instead the integrated object was plotted using nuclei type labels to permit examination of which profiled subpopulations of putamen oligodendrocyte nuclei might have greater transcriptional similarity to nuclei from substantia nigra. SNCA expression in healthy substantia nigra was also examined using feature plots to compare expression among samples.

### Statistics and Reproducibility

Putamen snRNA-seq data and custom analysis code used in this study will be made available upon publication. snRNA-seq data files for substantia nigra are downloadable online [3]. Statistical methods are presented in each of the above sections in the context of their use and interpretation. Supplemental files provide gene lists for Control oligodendrocyte cluster markers, differentially expressed genes, and modules identified by WGCNA. For enrichment testing, we used hypergeometric test using the R function phyper(). For comparison of cell cluster proportions versus disease state, we used X-squared test using the R function chisq.test() to compare total proportions between groups.

## DATA AND CODE AVAILABILITY

Putamen single-nucleus data files: Data to be made available online at time of publication.

Seurat package software: Install from CRAN (https://satijalab.org/seurat/install.html)

WGCNA package: Installation and documentation (https://horvath.genetics.ucla.edu/html/CoexpressionNetwork/Rpackages/WGCNA/)

NetworkAnalyst online interface (https://www.networkanalyst.ca/)

Enrichr: Computational systems biology interface (https://amp.pharm.mssm.edu/Enrichr/)

SN single-nucleus data files: https://www.nature.com/articles/s41467-020-17876-0 (data on NCBI: https://www.ncbi.nlm.nih.gov/geo/query/acc.cgi?acc=GSE140231)

R notebooks with custom analysis code will be made available online at time of publication.

## ACKNOWLEDGMENTS

We thank Dr. Srinivas Shankara for critical review and insightful feedback on this paper.

## AUTHOR CONTRIBUTIONS

All authors have each made substantial contributions to study concept, design, and implementation; drafting and critically revising the manuscript for important intellectual content; and provided final approval of the submitted manuscript version.

## Figure and Table Legends

**Supplementary Figure 1.**
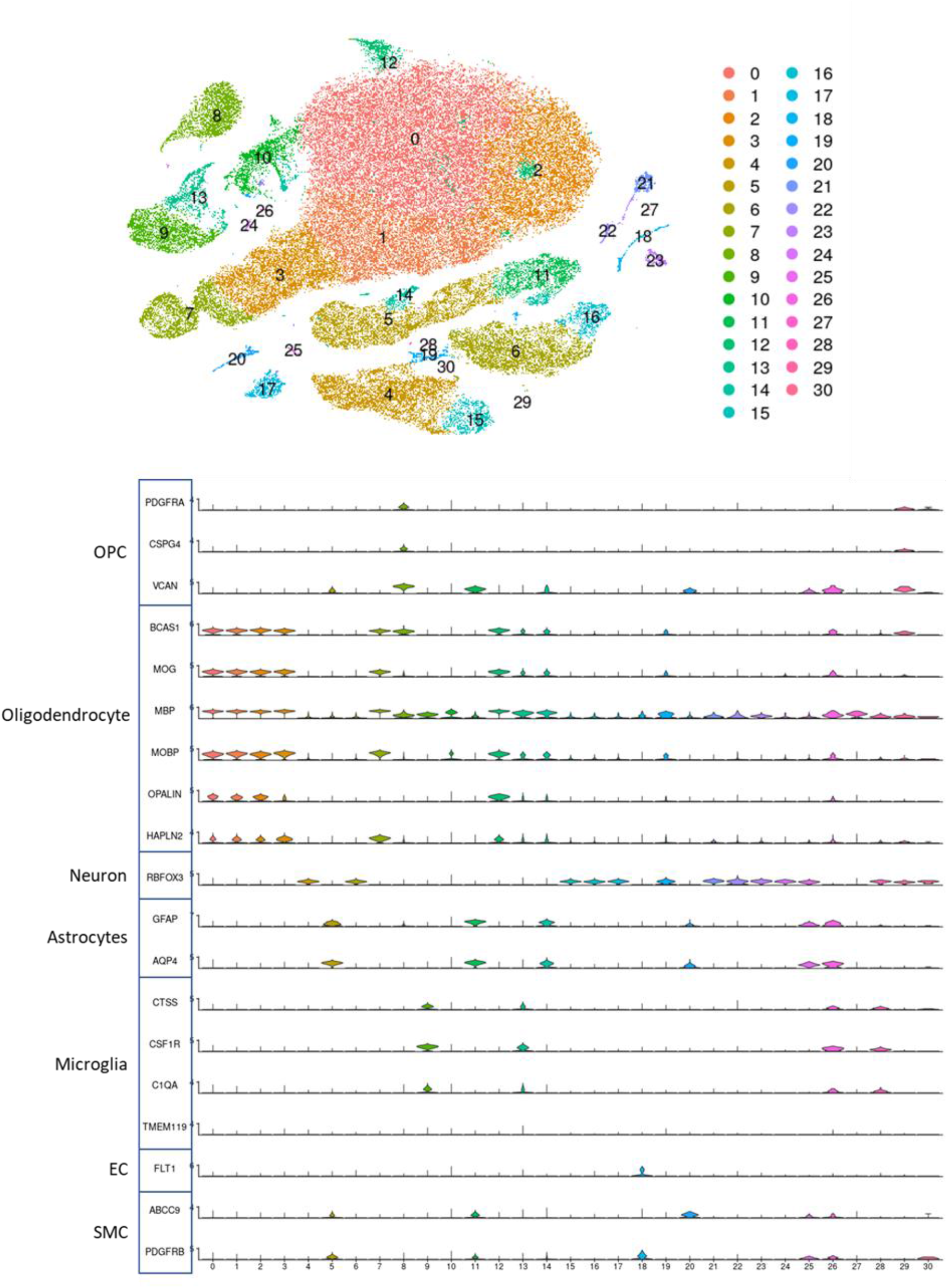
Broad cell types annotation for integrated PD, MSA, and Control samples.

**Supplementary Figure 2.**
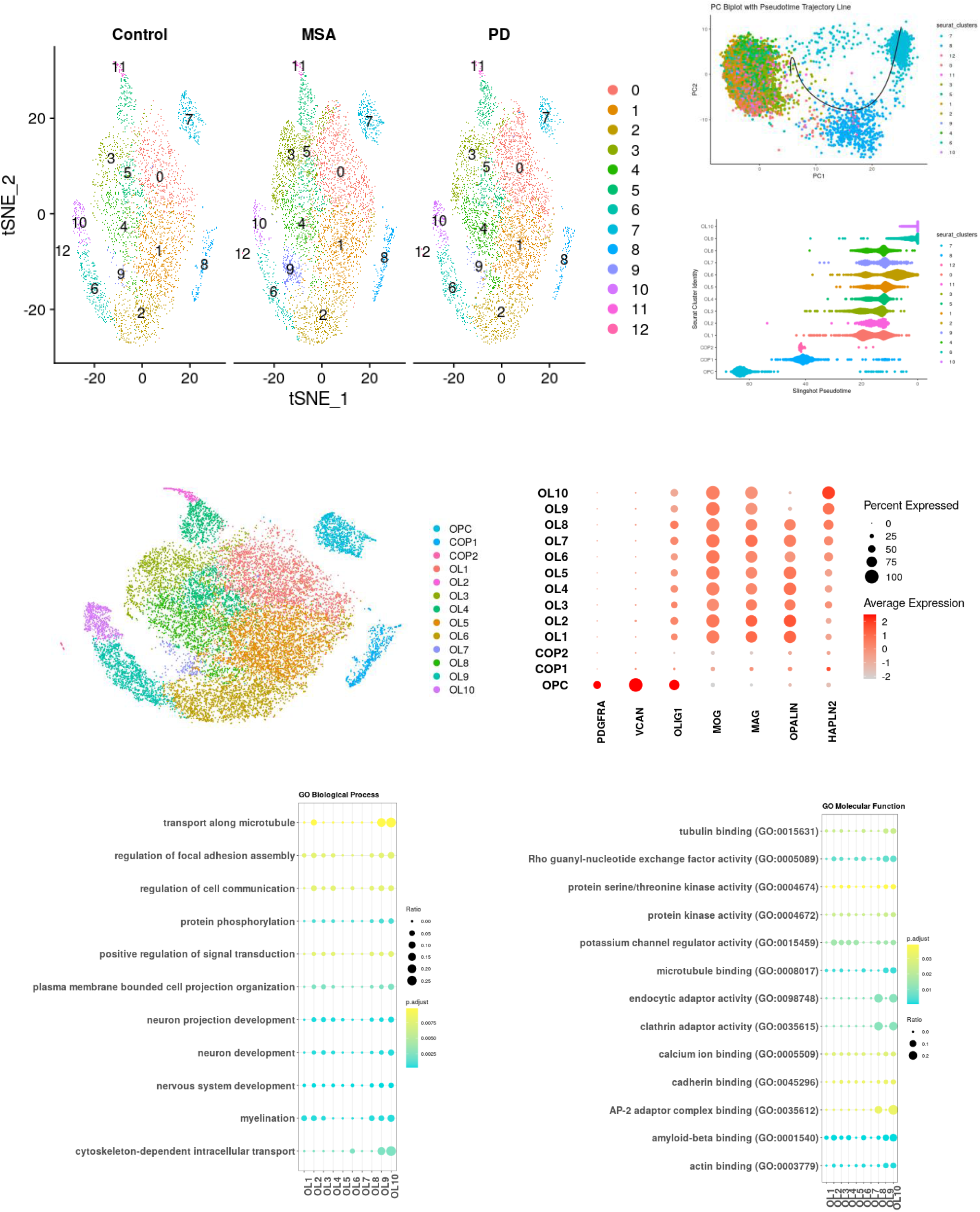
Control Oligodendrocytes. Clustered nuclei for CONTROL ordered by pseudotime and maturation marker expression. Groups with similar pseudotime and marker expression profiles combined for downstream analysis. Cluster assignments for downstream analysis are as follows: COP:[COP1,COP2]; OL1:[OL1,OL2]; OL2:[OL3,OL4];OL3[OL5,OL6]; OL4:[OL7,OL8]; OL5: [OL9,OL10].

**Supplementary Figure 3.**
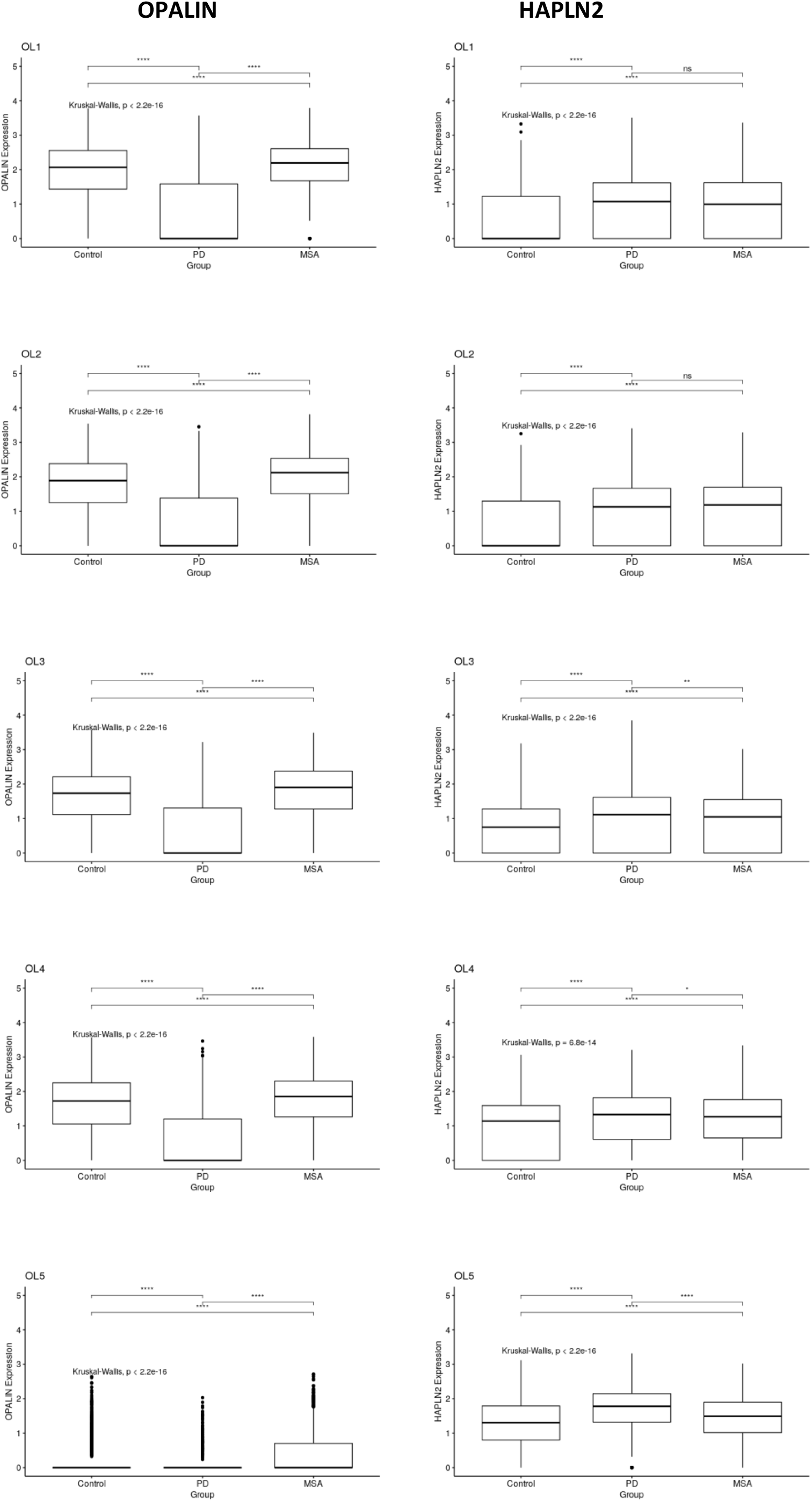
Gene expression significance tests for OPALIN and HAPLN2 by OL cluster.

**Supplementary Figure 4.**
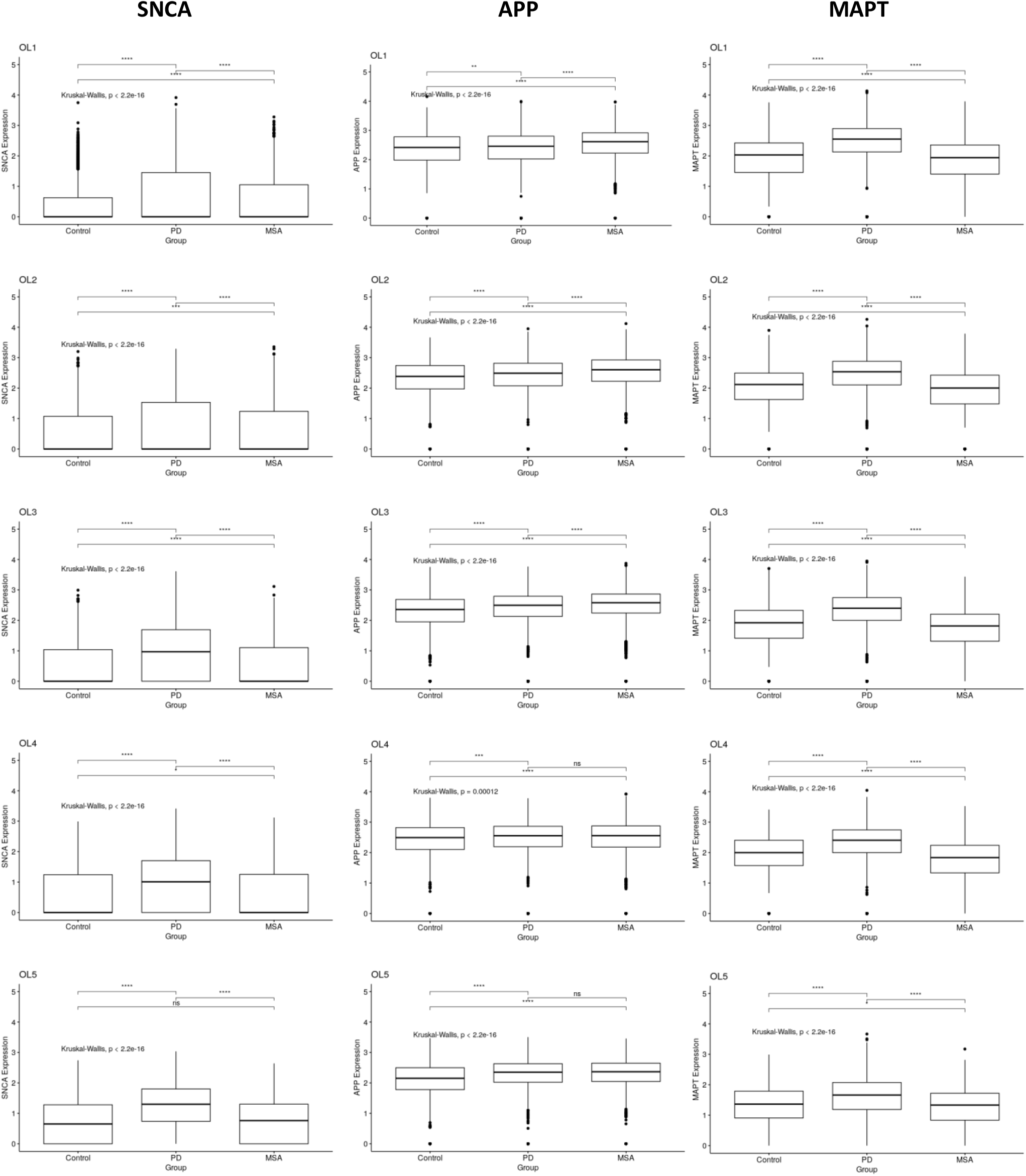
Gene expression significance tests for SNCA, APP, and MAPT by OL cluster.

**Supplementary Fig. 5.**
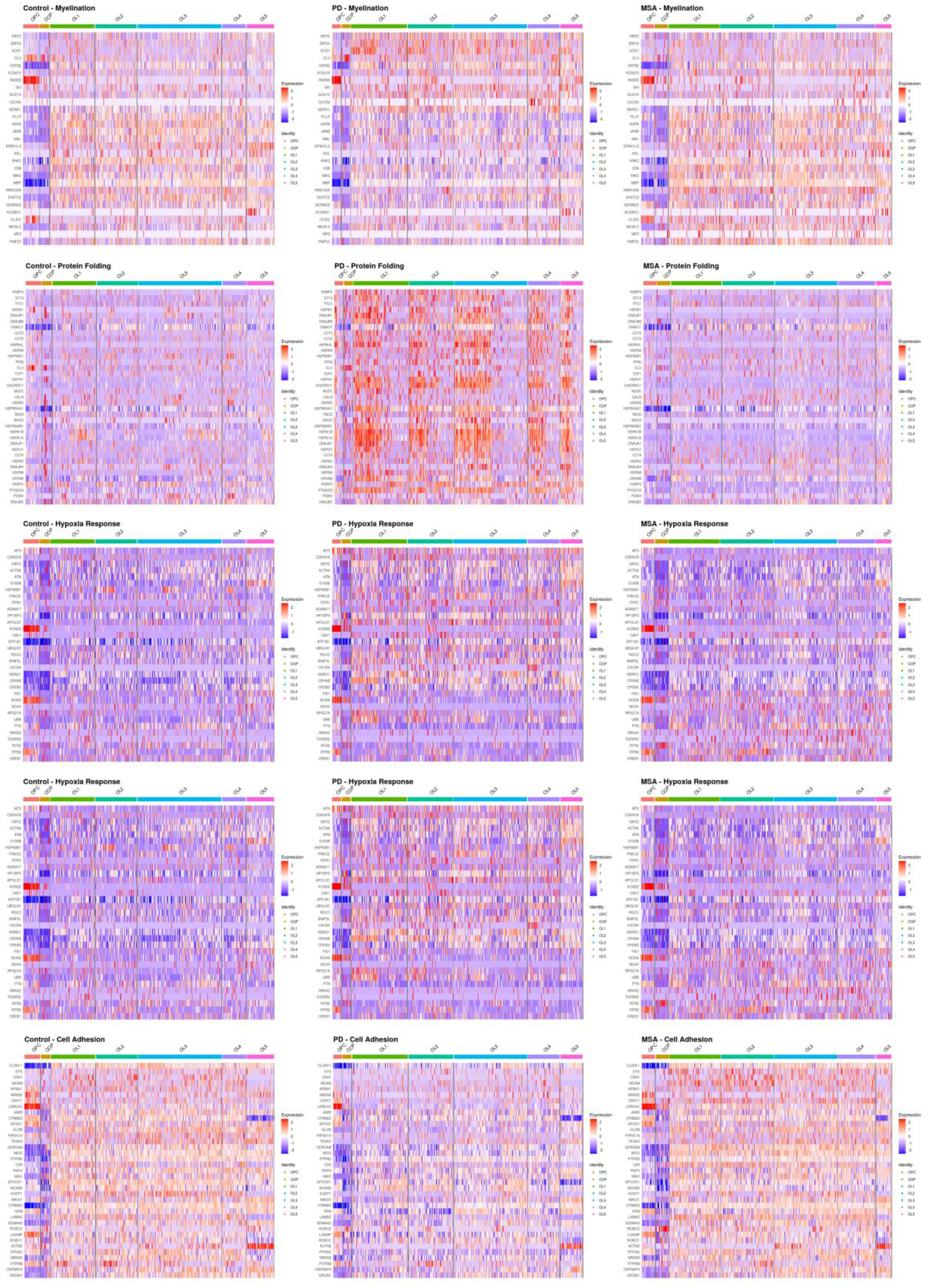
Expression by cluster and condition for selected enriched pathway by differentially expressed gene sets (integrated scaled data shown for −2 to 2 range, blue: downregulated, red: upregulated).

**Supplementary Fig. 6.**
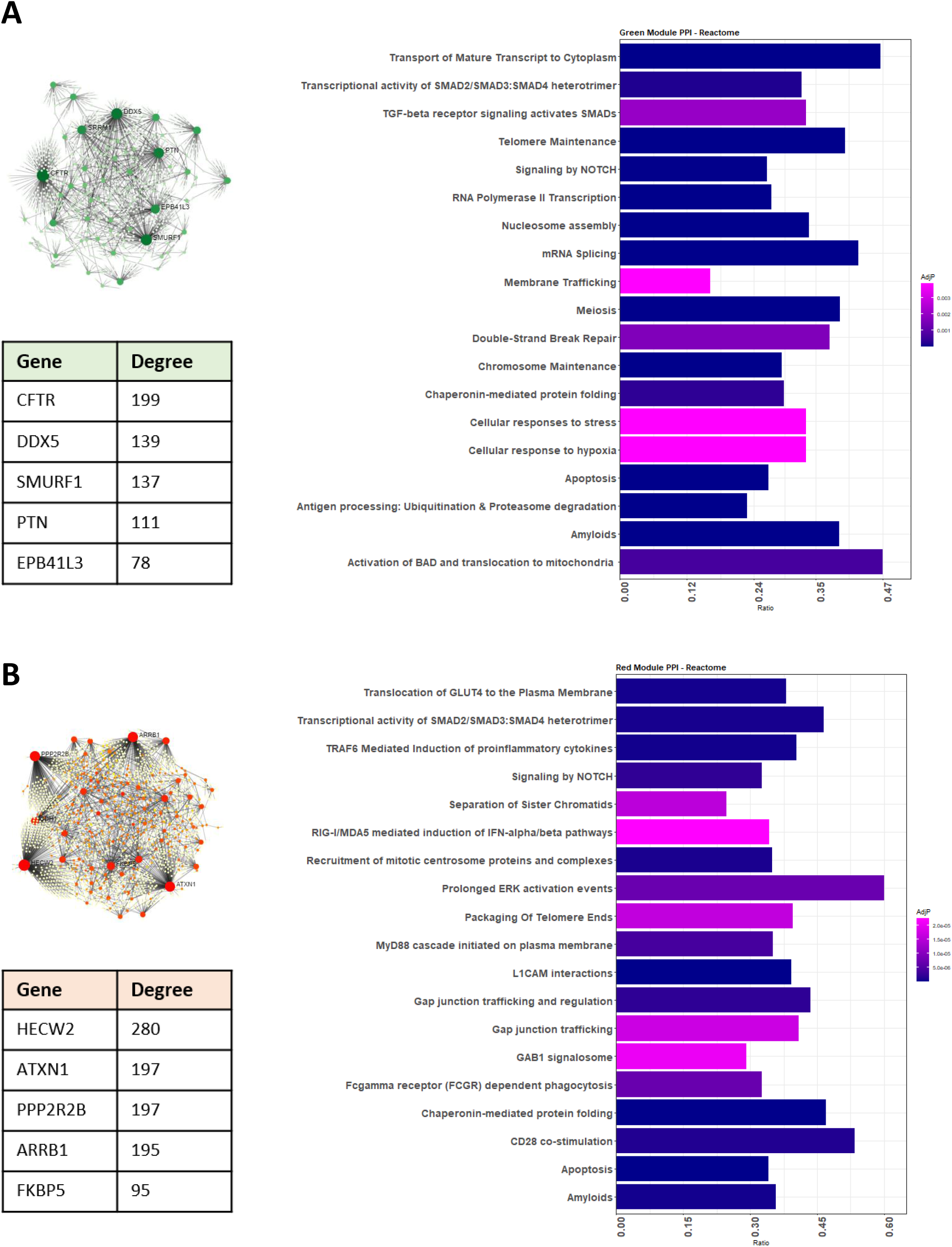
WGCNA. identifies co-expression modules significantly positively correlated with MSA and negatively correlated with PD. Reactome enrichment of PPI networks for these modules are shown: Green (A) and Red (B).

**Supplementary Fig. 7.**
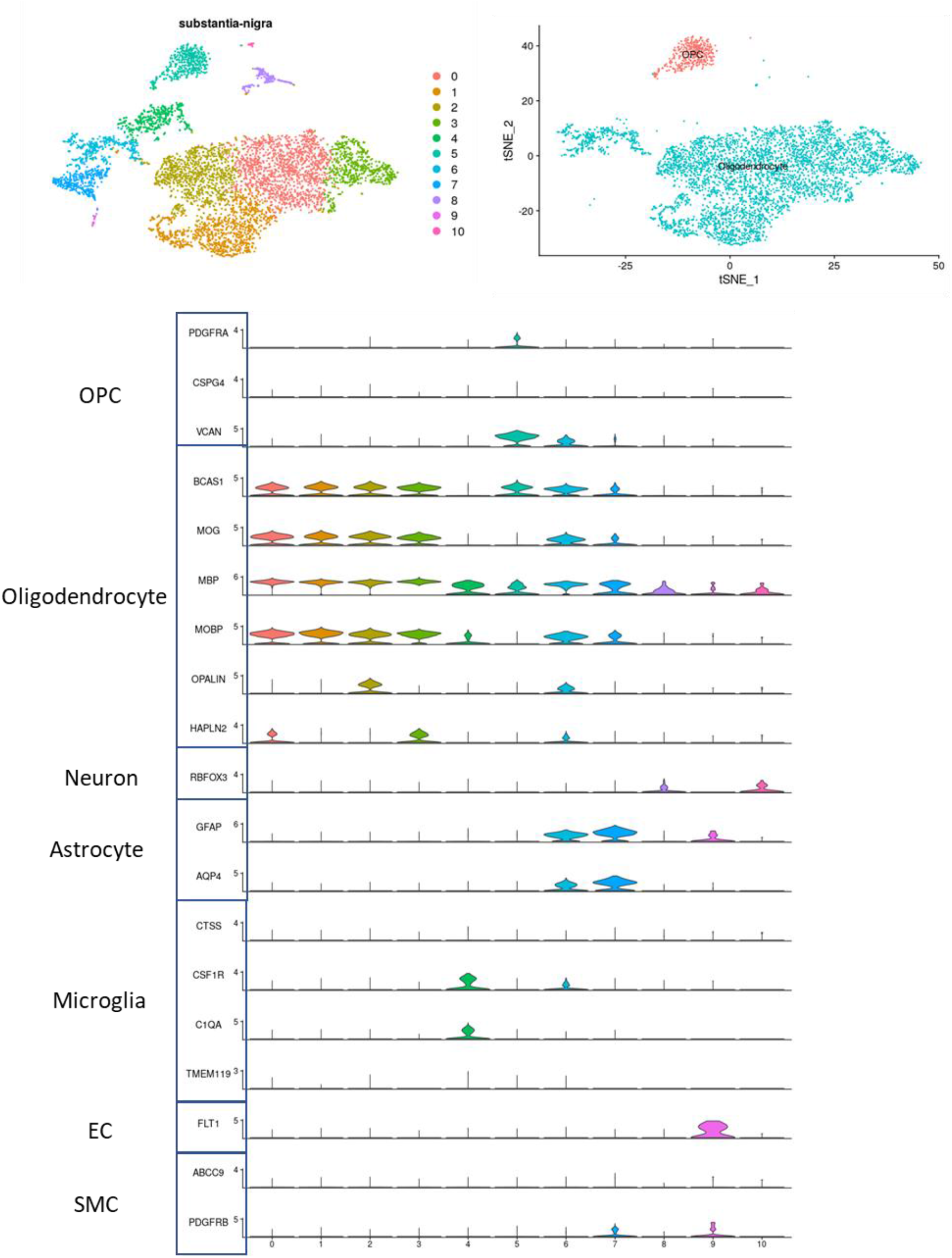
Broad cell types annotation for integrated substantia nigra samples.

**Supplement Table 1.** Sample Information

| ID | Group | Age | Sex | Nuclei | Post-QC Nuclei* | Post-QC nFeatures/cell |
| --- | --- | --- | --- | --- | --- | --- |
| 1 | CTR | 69 | M | 5044 | 4960 | 15147 |
| 2 | PD | 76 | M | 10762 | 10726 | 15217 |
| 3 | CTR | 88 | M | 7132 | 6895 | 15492 |
| 4 | PD | 77 | M | 12272 | 12012 | 15583 |
| 5 | MSA | 61 | M | 10213 | 10102 | 15541 |
| 7 | MSA | 57 | M | 11139 | 11047 | 15699 |
| 8 | CTR | 79 | M | 10496 | 10442 | 15523 |
| 9 | PD | 86 | M | 9624 | 9563 | 15252 |
| 10 | MSA | 77 | M | 11490 | 11339 | 15684 |

\*QC filters: nFeature\_RNA>200&<9000; percent.mt<5

**Counts of Nuclei Broad Types by Sample for Putamen**
| ID | Group | Oligodendrocyte | OPC | Neuron | Astrocyte | Microglia | SMC_EC |
| --- | --- | --- | --- | --- | --- | --- | --- |
| 1 | CTR | 3736 | 221 | 436 | 347 | 216 | 4 |
| 2 | PD | 9214 | 256 | 127 | 598 | 522 | 9 |
| 3 | CTR | 3288 | 464 | 1665 | 1121 | 299 | 58 |
| 4 | PD | 4358 | 255 | 3746 | 2540 | 962 | 151 |
| 5 | MSA | 7991 | 411 | 507 | 658 | 521 | 14 |
| 7 | MSA | 5875 | 552 | 3180 | 981 | 360 | 99 |
| 8 | CTR | 7595 | 412 | 1464 | 623 | 307 | 41 |
| 9 | PD | 6502 | 264 | 1011 | 1249 | 460 | 77 |
| 10 | MSA | 6743 | 261 | 2797 | 843 | 572 | 123 |

**Counts of Nuclei Oligodendrocyte Lineage Subtypes by Sample for Putamen**
| ID | Group | OPC | COP | OL1 | OL2 | OL3 | OL4 | OL5 |
| --- | --- | --- | --- | --- | --- | --- | --- | --- |
| 1 | CTR | 215 | 104 | 618 | 504 | 1585 | 205 | 722 |
| 2 | PD | 251 | 137 | 2294 | 1427 | 3104 | 1258 | 996 |
| 3 | CTR | 386 | 216 | 484 | 801 | 1118 | 535 | 135 |
| 4 | PD | 250 | 477 | 954 | 1151 | 1095 | 569 | 114 |
| 5 | MSA | 392 | 129 | 1660 | 1184 | 2999 | 1053 | 971 |
| 7 | MSA | 545 | 368 | 1500 | 1798 | 985 | 1020 | 204 |
| 8 | CTR | 406 | 296 | 1702 | 1337 | 2605 | 778 | 877 |
| 9 | PD | 257 | 138 | 1471 | 1223 | 2013 | 873 | 787 |
| 10 | MSA | 246 | 681 | 1342 | 1681 | 1593 | 1248 | 200 |

Counts of Oligodendrocyte Lineage Predicted Subtypes for Substantia Nigra
| ID | SN-ID | OPC | COP | OL1 | OL2 | OL3 | OL4 | OL5 |
| --- | --- | --- | --- | --- | --- | --- | --- | --- |
| 1 | N1B | 46 | 18 | 135 | 0 | 17 | 0 | 342 |
| 2 | N2B | 134 | 15 | 207 | 0 | 283 | 0 | 544 |
| 3 | N4B | 88 | 57 | 61 | 0 | 22 | 0 | 98 |
| 4 | N5B | 82 | 19 | 9 | 0 | 36 | 0 | 200 |
| 5 | N3 | 61 | 78 | 927 | 0 | 198 | 0 | 171 |
| 6 | N4 | 126 | 34 | 79 | 0 | 25 | 0 | 168 |
| 7 | N5 | 18 | 8 | 78 | 0 | 165 | 0 | 336 |

**Supplement Table 2.** Markers for Broad Cell Types Annotation

| Cell Type | Marker List |
| --- | --- |
| Oligodendrocyte | BCAS1, MOG, MBP, MOBP, OPALIN, OLIG1, HAPLN2 |
| OPC | PDGFRA, CSPG4, VCAN |
| Neuron | RBFOX3 |
| Astrocyte | GFAP, AQP4, GJA1 |
| Microglia | CTSS, CSF1R, C1QA, TMEM119, PTPRC, |
| Endothelial Cells | FLT1 |
| Smooth Muscle Cells | ABCC9, PDGFRB |

**Supplement Table 3.**
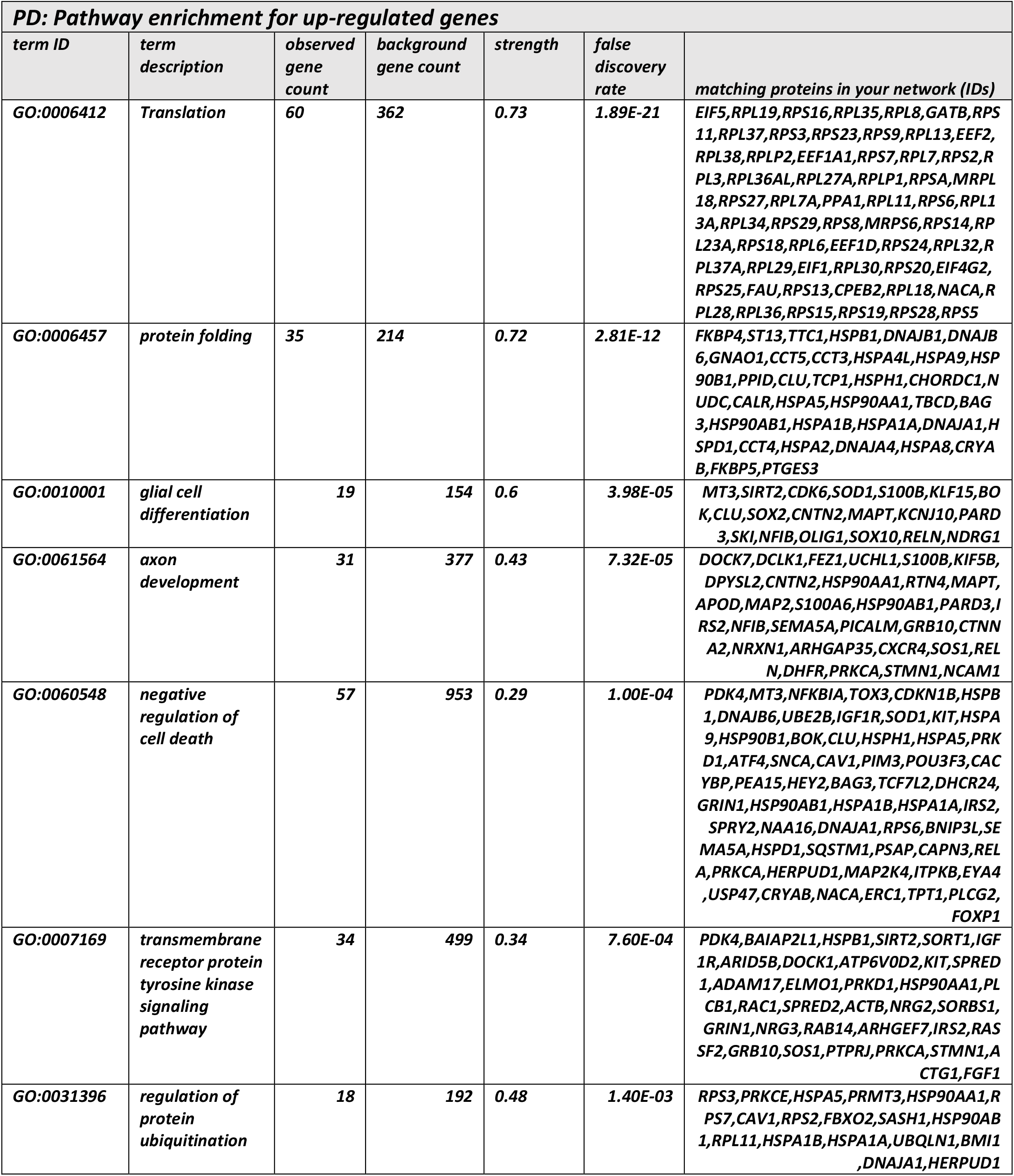
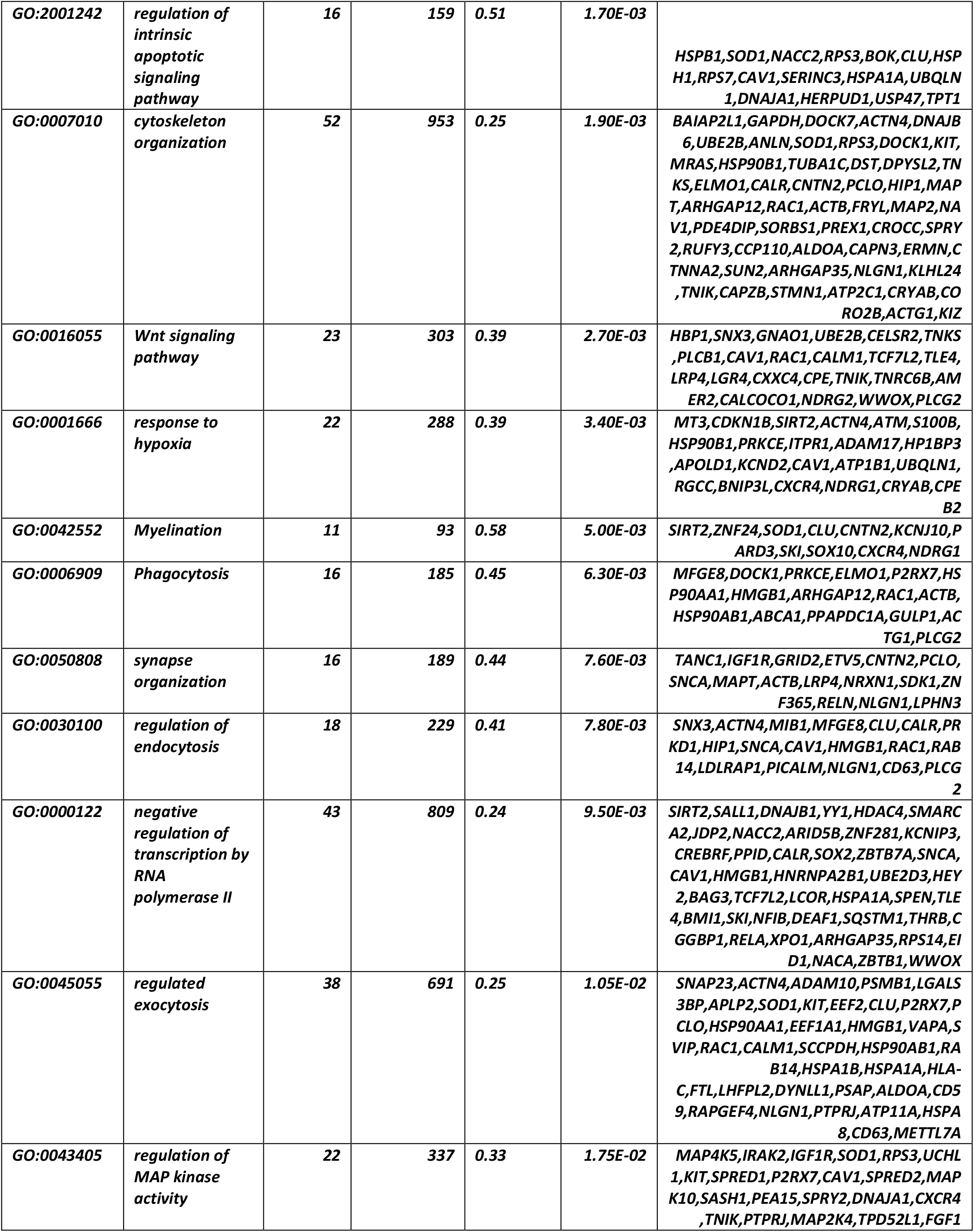
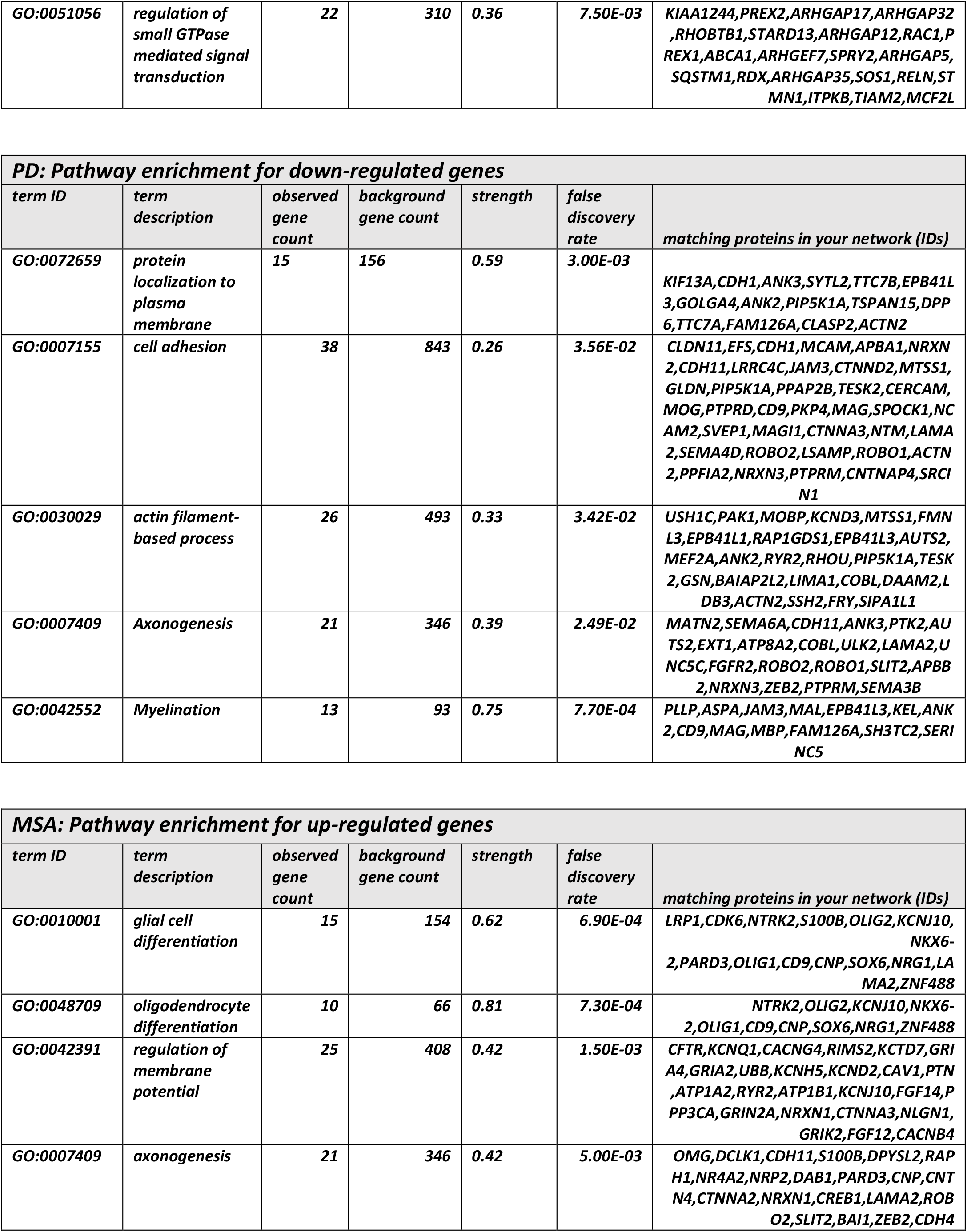
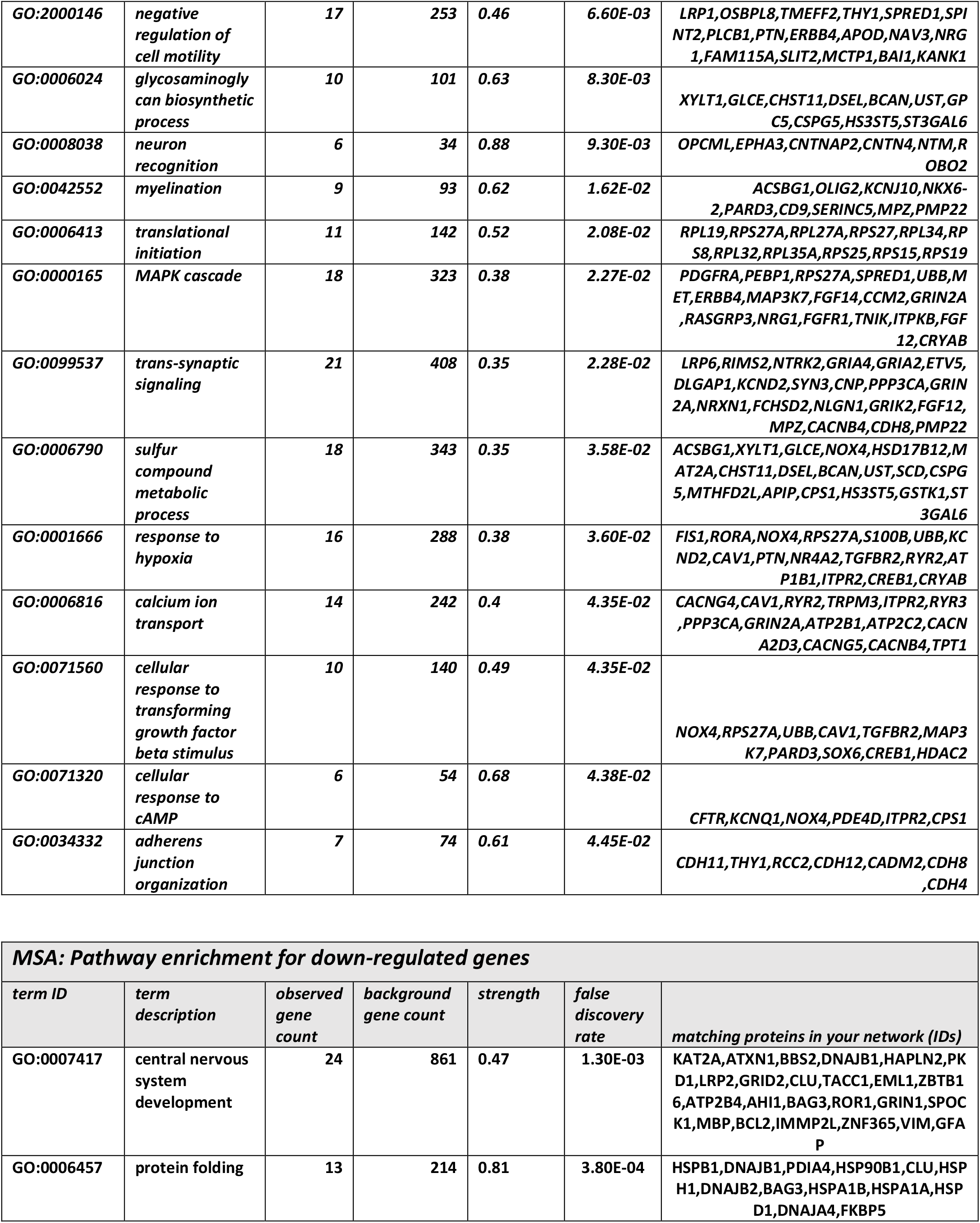
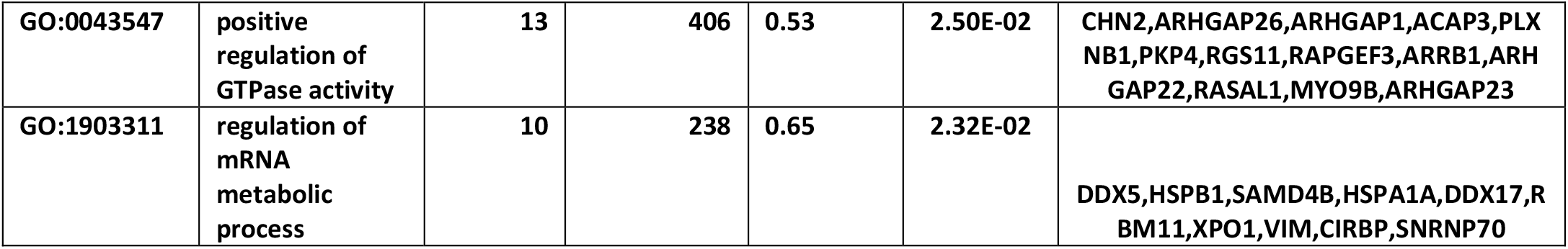
Pathway Enrichment for Differentially Expressed Genes in PD and MSA (combined cluster results).

